# Not seeing the forest for the trees: Combination of path integration and landmark cues in human virtual navigation

**DOI:** 10.1101/2023.10.25.563902

**Authors:** Jonas Scherer, Martin M. Müller, Patrick Unterbrink, Sina Meier, Martin Egelhaaf, Olivier J. N. Bertrand, Norbert Boeddeker

## Abstract

**Introduction:** In order to successfully move from place to place, our brain often combines sensory inputs from various sources by dynamically weighting spatial cues according to their reliability and relevance for a given task. Two of the most important cues in navigation are the spatial arrangement of landmarks in the environment, and the continuous path integration of travelled distances and changes in direction. Several studies have shown that Bayesian integration of cues provides a good explanation for navigation in environments dominated by small numbers of easily identifiable landmarks. However, it remains largely unclear how cues are combined in more complex environments.

**Methods:** To investigate how humans process and combine landmarks and path integration in complex environments, we conducted a series of triangle completion experiments in virtual reality, in which we varied the number of landmarks from an open steppe to a dense forest, thus going beyond the spatially simple environments that have been studied in the past. We analysed spatial behaviour at both the population and individual level with linear regression models and developed a computational model, based on maximum likelihood estimation (MLE), to infer the underlying combination of cues.

**Results:** Overall homing performance was optimal in an environment containing three landmarks arranged around the goal location. With more than three landmarks, individual differences between participants in the use of cues are striking. For some, the addition of landmarks does not worsen their performance, whereas for others it seems to impair their use of landmark information.

**Discussion:** It appears that navigation success in complex environments depends on the ability to identify the correct clearing around the goal location, suggesting that some participants may not be able to see the forest for the trees.

## 1 INTRODUCTION

Spatial navigation is one of the most crucial behavioural competencies of many animals, including humans. The cognitive processes underlying spatial navigation abilities have been the focus of much research, with two major sources of spatial information being identified as the basis of spatial learning: landmarks (Chen et al., 2017; Zhao and Warren, 2015b; Mallot and Lancier, 2018; Jetzschke et al., 2017) and path integration (PI) (Jetzschke et al., 2016; Harootonian et al., 2022; Chrastil and Warren, 2021). Both are used by non-human and human navigators alike (Etienne and Jeffery, 2004). The use of landmarks most often refers to the usage of external environmental features to identify and locate places of interest in a navigator’s surroundings (Zhao and Warren, 2015b; Jetzschke et al., 2017; Walter et al., 2022). Landmarks can be used for guidance towards a goal (”turn left at the gas station”) as well as for self-localisation (”I see the Eiffel Tower, so I must be in Paris”). Path integration (PI), or “dead reckoning”, refers to the continuous integration of self-motion information (translation and rotation, gathered for example from visual and proprioceptive cues) (Wiener and Mallot, 2006; Chrastil et al., 2019) for the purpose of navigation.

### 1.1 Cue integration in homing

One of the most frequent and important spatial tasks navigators have to solve for any navigator is returning to a previously visited location, like an important food source or their home, hence called “homing”. Combining different sources of information (cues) can be a crucial strategy to improve success in homing, since the information provided by each individual cue is limited and noisy. PI accumulates errors, potentially decreasing its usefulness over time, while landmarks might be visible only from some part of the environment or be hard to identify in visually ambiguous situations (Hoinville and Wehner, 2018). When combining cues, the relative importance of each depends on a variety of factors, including availability, task context, environmental structure, and cue reliability, that all can be represented well in a Bayesian framework of optimality (Ernst and Banks, 2002; Alais and Burr, 2019). In such a framework, each source of information is weighted based on its perceived reliability and can be combined with prior knowledge about certain features of the environment (Zhao and Warren, 2015b; Chen et al., 2017; McNamara and Chen, 2022; Roy et al., 2023). In navigation research the application of the Bayesian framework predicts that the combined estimate of multiple noisy sources of information will be more precise (lower standard deviation) (Chen et al., 2017), while its overall accuracy (central tendency of magnitude of error) will depend most strongly on the accuracy of the estimate derived from the most reliable cue. Together, these two error measures form the basis of several experimental studies, which validate the prediction of optimal cue integration models and show that humans mostly combine cues in a statistically optimal manner, but are also prone to systematic misjudgements under some conditions (Zhao and Warren, 2015b; Jetzschke et al., 2017; Nardini et al., 2008; Zhao and Warren, 2015b; Chen et al., 2017; Sjolund et al., 2018; Kessler et al., 2022, preprint). One effect of landmark ambiguity was demonstrated by Jetzschke et al. (2017), who could show that homing performance, i.e. the ability to return back to one’s starting position, improved when human navigators were able to make use of multiple local landmarks close to their goal location. However, cue integration also made participants susceptible to being misled, with one landmark being covertly displaced and participants not perceiving the mismatch they headed towards an intermediate location. Several other studies show that when a mismatch between cues is detected, navigators rely on only one of the spatial cues available to them, either in form of cue competition or cue alternation (Zhao and Warren, 2018; Sjolund et al., 2018; Harootonian et al., 2022). First, in cue competition, available cues compete against each other in decision making. The dynamics of such competition are matter of ongoing debate (Zhao and Warren, 2018; Harootonian et al., 2022; Kessler et al., 2022, preprint), but it could be shown that when, for example, PI and landmark cues are put in conflict by rotating them against each other, the two cues compete and participants choose landmark cues for homing when the rotational conflict is small, but ignore them when the conflict is large (Zhao and Warren, 2015b; Sjolund et al., 2018). Second, in cue alternation, navigators alternate between cues from trial to trial, with the probability of choosing either cue being based on perceived cue reliability, meaning that more reliable cues are used more often. This resembles “probability matching”, a known decision making strategy in perceptual and cognitive tasks (Wozny et al., 2010). In order to disentangle the combination of information from landmarks and PI in human navigation, homing tasks have received much attention in the spatial cognition literature (e.g. Jetzschke et al. (2017); Chen et al. (2017); Zhao and Warren (2015b); Roy et al. (2023); Widdowson and Wang (2022)). One of the most prominent homing tasks is the triangle completion task (e.g. Loomis et al. (1993); Chen et al. (2017); Harootonian et al. (2020); Zhao and Warren (2015b)). In a triangle completion task, the participant is actively or passively guided along two sides of a triangle, and at the end of this outbound path asked to either actively return to, or point at the starting location (Kearns et al., 2002; Glasauer et al., 2002; Wolbers et al., 2007). Comparing systematic errors in different conditions (e.g. with or without landmarks (Zhao and Warren, 2015b), longer or shorter paths (Harootonian et al., 2020), or different turning angles (Wiener and Mallot, 2006; Harootonian et al., 2020), or for healthy individuals vs. those suffering from vestibular disorders (Xie et al., 2017)) allows to draw conclusions about how these factors might have influenced behaviour. Manipulating the reliability of cues and observing their influence on performance in different environments in a triangle completion task also allows to investigate Bayesian cue integration models, as cue weighting in these models depends on cue reliability. Thus, among other factors, homing accuracy and precision seems to be dependent on the specific experimental conditions and the environment used, as they directly impact reliability of PI and landmark cues.

### 1.2 Influence of environmental complexity

Recent experimental and theoretical studies have started to explore the relation between the systematic errors observed in cue combination and the characteristics of the environment within which a behaviour was observed. It was shown that navigators can re-assess the reliability of spatial information extracted from an environment based on changes in its spatial structureRoy et al. (2023). Zhao and Warren (2015a) reported that participants often misjudge the reliability of visual cues in a homing task and even continue to use landmark cues for visual guidance when objects were displaced and put in conflict with the internal PI of the participants. However, the studies by Roy et al. (2023) and Zhao and Warren (2015a) also report that if participants successfully identify objects as spatially unstable (i.e. appearing at different locations throughout the experiment), this knowledge is quickly incorporated into the participant’s navigation strategy, resulting in reduced trust in landmark-based guidance and increased reliance on PI. These effects of environmental context also seem to be expressed differently in different people, leading to individual differences in short-term learning (Glasauer and Shi, 2022) and the relative reliance on different spatial cues (Zanchi et al., 2022). The findings by Jetzschke et al. (2017), Roy et al. (2023), and Zanchi et al. (2022) lead us to an important conclusion: If the use and combination of spatial cues for navigation depends on a) the makeup of the environment and b) the nature of the task at hand, it becomes necessary to more explicitly consider these factors in designing our experiments, if we want to deepen our understanding of the mechanisms that govern (human) spatial cognition. In other words, for a given task, we can categorise different environments based on how easy it will be to solve that type of task in the respective environment. This in turn will depend on different aspects of the makeup of that environment, such as the number, distribution, and distinguishability of spatial features in it. We will call the totality of these features the “complexity” of that environment. Building on this, we can ask how changing specific features contributing to the complexity of an environment will affect the ability of human navigators to solve a spatial task and also which strategies they might employ to do it. As an example, consider a meadow with a single tree on it. If we need to memorise and return to a location close to that tree, we can assume that the tree will serve as a useful landmark, providing us with information about the relative distance between the goal location and itself. Now consider we add a large number of similar-looking trees to this meadow. Geometrically, the information provided by the original tree did not change. However, when navigating in this new, more complex environment we might on the one hand make use of new, emergent cues, such as spatial configurations of groups of trees. On the other hand, using any given tree or configuration of trees at all might be limited to the degree we can identify the tree(s) in question. This illustrates how cue use can be closely tied to the complexity of a spatial environment. However, so far, most experimental studies concerned with cue combination and cue preference have been conducted in relatively simple environmental settings and have not systematically addressed the question of visual complexity and its influence on homing.

### 1.3 How does homing performance of human navigators depend on the visual complexity of the environment?

We aim to answer the above question by systematically varying the degree of spatial complexity in a spatial homing task. Our study is guided by two hypotheses when comparing homing behaviour in environments with different numbers of ambiguous landmark objects. First, if navigators are able to identify individual objects (e.g. based on the spatial configuration of object groups or by disambiguating landmark positions based on PI information), homing errors should decrease as more objects are added, up to a point where the goal location can be completely constrained based on the spatial information derived from landmark cues. This is the situation described by Jetzschke et al. (2017), who could show that homing performance was best when participants could fully triangulate the goal from the three present landmark cues. Second, if objects become ambiguous to the navigator, as could be the case if more objects are added beyond what is needed to geometrically constrain the goal location, homing performance should decrease as more and more ambiguous objects in the environment increase the chances for the navigator to be lead astray. This situation has not been investigated yet and is challenging the cue integration model proposed by Jetzschke et al. (2017). Considering the increasing cognitive load necessary to differentiate a rising number of landmarks in complex environments, memorise the distance to these cues, or use emerging landmark configurations, we do not expect the proposed model to well depict cue integration in complex environments. Thus, the usefulness of landmark cues for homing might be smaller in more complex environments and in extreme cases, the navigator might rely solely on PI to find their goal. However, when taking into account results like those of Zanchi et al. (2022), who report large inter-individual differences in cue use even in relatively simple environments, we expect that reliance on the different cues should vary between participants in more complex environments, maybe even to a greater degree than in simple environments. We therefore expect that homing performance is subject to inter-individual variability and will explicitly include this factor in our analysis.

To systematically assess the effects of environmental complexity on homing behaviour, we have performed a virtual reality (VR) experiment on desktop computers. The use of VR tools is of special interest for this type of experiment since it allows for the real-time presentation and manipulation of complex, interactive environments. We made use of these capabilities to design a triangle completion task with active return to the home location. Crucially, the environment differed between experimental conditions, containing different numbers of visually ambiguous trees surrounding the goal location, creating different degrees of clutter. We take these different degrees of clutter to be our proxy for the abstract idea of “environmental complexity” in this study. This experimental paradigm allows us to target the question of how homing performance and spatial cue integration depend on the visual complexity of an environment by systematically introducing additional, ambiguous objects.

## 2 MATERIALS AND METHODS

### 2.1 Task description

This study is using a triangle completion task, in which the participant is guided away from a goal location along two legs of a triangle and then asked to return to their goal location, thus trying to complete the triangle as accurately as possible. By quantifying the errors made by participants, this task assesses the participant’s homing performance. We tested participants in triangle completion tasks in different conditions that differed in the number of available tree objects in the surroundings. On the one hand, this study aims to verify and build upon the findings by Jetzschke et al. (2017) for spatially simple environments. For verification in our visual VR paradigm we test participants in corresponding environments with zero, one, two, and three identical, rotationally symmetric trees to present different degrees of visual clutter and thus different degrees of visual ambiguity of the possible landmark cues. On the other hand, we aim to challenge the proposed cue integration model by asking our participants to also navigate in intermediate and high degrees of visual clutter in environments with ten and 99 visually ambiguous landmarks. The condition with zero objects was included to provide a baseline of performance for a situation in which only PI information was available for navigation. Conditions with higher object counts always contained the objects of conditions with lower object counts as well (Fig. 1).

**Figure 1.**
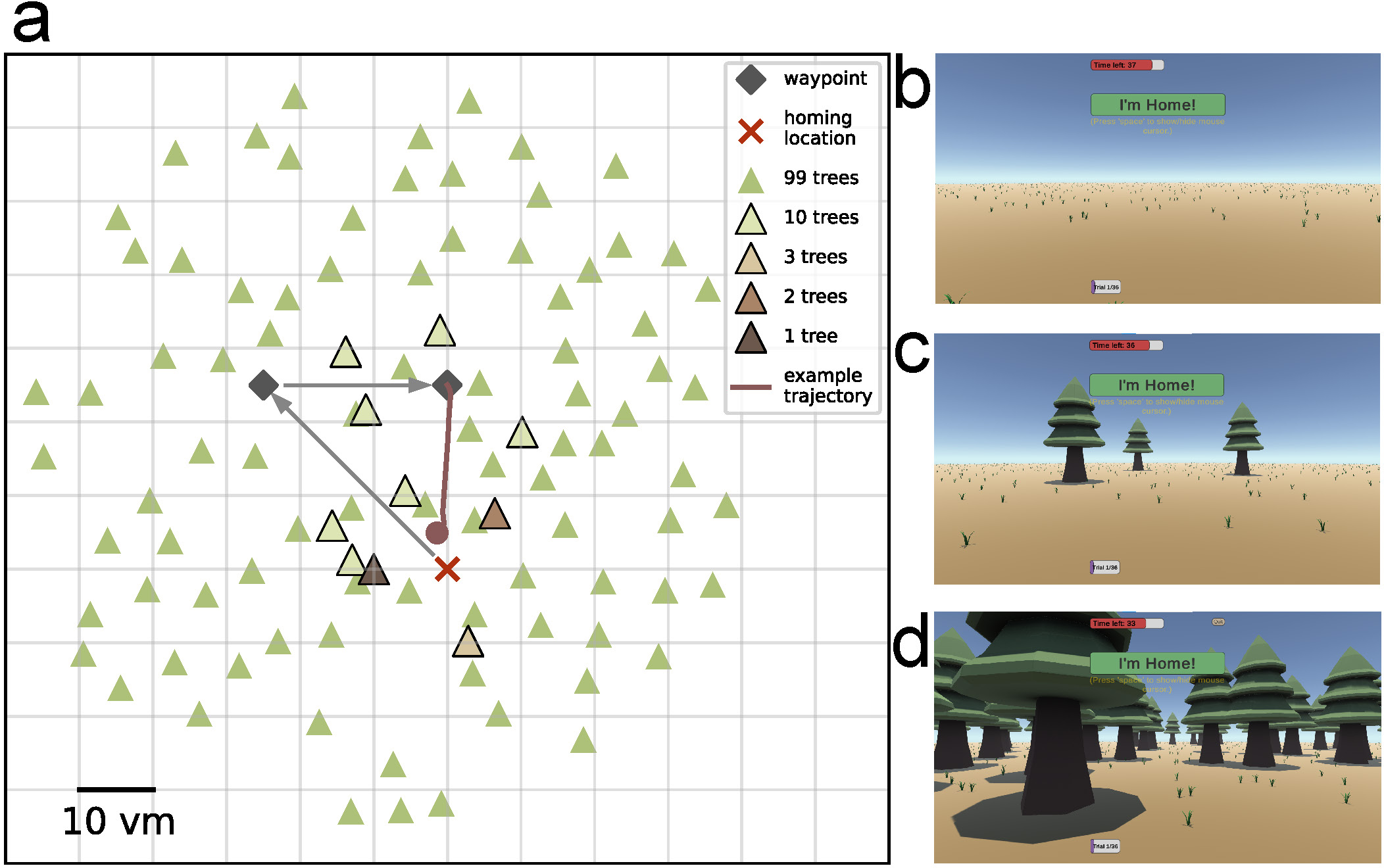
Triangle completion task and environmental complexities across conditions. (a) Participants are guided away from a goal location along two sides of a triangle. The goal location is indicated in our VR environment by a campfire, and waypoints by piles of firewood, that participants are asked to collect because their campfire went out. Waypoint locations are not changed and participants in all repetitions and conditions walk the same outbound path and are asked to return to their goal location, the campfire, at the end (example trajectory). Environmental complexity is varied by altering the number of available objects between zero, one, two, three, ten, and 99. The conditions with higher numbers of objects contain also the objects from conditions with a lower number of objects. Scale is given in virtual meters (*vm*), that correspond to actual meters in the VR. Subfigures b, c, and d show screenshots of the first-person view from the VR for conditions with zero, three, and 99 objects, respectively, at the second waypoint in the direction of the goal location. UI elements shown in the screenshots from top to bottom are: 1) a red timeout bar, indicating the time left for the trial; 2) an ”I’m Home!’ button, to log in the final position; 3) the written instruction “Press ‘space’ to show/hide mouse cursor” below; 4) and a purple bar indicating the current trial within the session.

In our study, all trees look exactly the same, to focus on the overarching effect of environmental complexity in the form of clutter and not on the specifics of landmark salience or individual object identification capabilities. Thus, trees are designed to be rotationally symmetric to not provide additional direction cues. This way, we have created a challenging navigation task, akin to finding one’s way through a dense spruce forest where single trees are hard to distinguish from one another and landmark-based guidance needs to rely on cognitive heuristics such as emerging geometric features. Therefore, while the individual trees might serve as landmarks, we cannot directly determine how participants derive spatial information from the trees and will be referring to them only as “objects” throughout the text.

### 2.2 Experimental procedure

25 healthy participants (aged 19-30 years, 9 self-identified as female, 16 self-identified as male) took part in the experiment. Participants were informed about the general procedure of the experiment but not about its specific purpose or the nature of the different conditions. All participants gave their written informed consent for participating and received monetary compensation (7 Euros per hour). The experiments were approved by the Bielefeld University Ethics Committee and conducted in accordance with the guidelines of the Deutsche Gesellschaft für Psychologie e.V. (DGPs), which correspond to the guidelines of the American Psychological Association (APA). The experiment was carried out over six sessions (one training + five main sessions), split evenly over two experimental days, which were not more than 1 calendar day apart. The experiment was provided in the form of a stand-alone Windows application, created using the virtual navigation toolbox for the Unity3D game engine and was executed remotely on participants’ desktop home computers (Müller et al., 2023). A supervisor was present to instruct and assist participants according to a standardised protocol during the initial training session via video call. After successful training, participants were able to complete the remaining experimental sessions without further supervision. However, participants were instructed to take regular breaks in between experimental sessions of at least 5-10 minutes. Each main session consisted of six repetitions of each of the six experimental conditions (number of objects) in a randomised order, leading to a total of 36 trials per session and adding up to n=24 repetitions per condition overall for each participant over the whole experiment. Each session took roughly 30-45 minutes to complete, leading to a total experiment duration of 3-4.5 hours per participant.

### 2.3 Trial structure

During a single trial of the experiment, the participant was first guided to the goal location, marked by a stylised campfire in the virtual environment. The avatar was controlled from a first-person view using the up/down and the left/right arrow keys, respectively, for translation and rotation. After reaching the campfire goal location, they were tasked with collecting pieces of firewood, with the wood appearing in sequence at the two corners of the first two legs of the triangle path, being 35.4 and 25 *vm* long with an inner angle of 45*^◦^* (see Fig. 1). This was termed the outbound phase of the trial. During this initial phase, a floating arrow in front of participants indicated the correct direction towards the next pile of wood, but the participants were otherwise free to move as they liked. It was ensured that no trees were placed directly onto the outbound, or direct return path, so that participants did not have to maneuver around them. However, we also placed trees at the end of these alleys to ensure similar visual clutter in all directions. To ensure an overall straight path along the legs of the outbound path, the participant was confronted with a time limit (45 seconds) to collect all the firewood. The remaining time was always visible in the form of a countdown bar at the top of the screen. Once the participant had completed the outbound phase, they were asked to return to the campfire (i.e. the goal location), once again while being presented with a countdown timer (45 seconds). Neither the piles of wood nor the campfire were visible during the return, to prevent their use as spatial cues. Identical grass tufts on the ground were randomly arranged for each trial to enable visual estimation of self-motion by optic flow. A sun-like light source provided global, uni-directional illumination from directly above so that no directional information could be derived from shadows in the environment. The avatar moved at 2.5 virtual meters per second. During each trial, the position and rotation of the player’s avatar in the virtual environment were recorded for every rendered frame. All data collected from the participants and during the experiment was handled according to the General Data Protection Regulation (GDPR) of the European Union.

### 2.4 Analysis

All analyses, except for the linear regression analysis which was done in custom R (R Core Team, 2022) scripts in version 4.1.3, were conducted in custom Python (Van Rossum and Drake (2009))^1^ scripts in version 3.9.12.

#### 2.4.1 Error measure

The primary error measure for our analysis is the position error, which is the Euclidean distance between the intended goal location and the endpoint of the actually walked trajectory. We calculated median position errors for each participant and condition as our accuracy measure, and standard deviation of position error as our precision measure (the lower the standard deviation the higher the precision).

#### 2.4.2 Linear mixed effects models

Regression models are commonly used to analyse the relationship between two or more variables to make predictions and identify trends in data. Especially powerful are linear mixed effects (LME) models, that include fixed and random effects. Fixed effects refer to a study’s factors of interest, while random effects describe randomly sampled factors, such as participant identity. Taking both effect types into account, results in more accurate and reliable estimates of complex relationships in the data. Based on the distribution of position errors across the conditions, we fit LME models to identify the underlying mechanisms that lead to the observed changes in performance. We aim for a model to quantify the influence of number of objects and inter-individual variability on position error accuracy and precision. LME models assume a normal distribution of residuals of the response variable, which in our case is the median position error (accuracy), or its standard deviation (precision). Both the accuracy and precision measures are zero-bounded, and thus their residuals in a regression analysis cannot be expected to be normally distributed. Thus, all LME models were calculated using the log-transformed position error and its log-transformed standard deviation respectively. We fit a model predicting the log-transformed position error (and separately its standard deviation) from the number of objects as an ordinal fixed effect and the participant identity as a categorical random effect. We present two approaches to justify splitting the dataset into two separate effects, an effect of low clutter and an effect of high clutter, one based on a quadratic regression and one based on forward difference contrast coding (UCLA Statistical Consulting Group (2021))^2^. We quantify both effects by comparing differences in median position error and in its standard deviation between the respective conditions. Additionally, to understand how precision and accuracy relate to each other, we correlated them separately for each condition, and display results of exemplary participants performing at different degrees of accuracy and precision. In our LME model analysis, we observe a small trend of reduced position error over sessions (see Results “Disentangling the effect of different degrees of clutter” and supplement “S2 Session order effect”) that, however, only slightly improves the regression model fit. Since order effects are not a main focus of this study, we decide to average position errors for each participant within each condition (number of objects) using its median. Nevertheless, we also provide a secondary analysis using a LME model on single-trial data (see section “Data Availability Statement” for the supplemental notebooks). The alternative analysis reproduces the qualitative results presented in the main text.

#### 2.4.3 Maximum Likelihood Estimation modelling

Maximum likelihood estimation (MLE) is an important statistical tool for the analysis of navigation strategies (Chen et al., 2017). MLE assumes that navigators combine information from different sources in a way that maximises the likelihood of the observed experimental behaviour of a participant or population. To employ MLE as a tool, one must first formulate a mathematical model which describes the way the information sources in question could be used. MLE then seeks to find the set of parameters that best explain the observed data given the predefined model. Then, in a second step, one can compare the overall likelihood of different model variants, which are formal ways of describing qualitatively different uses of spatial information. This procedure can provide quantitative evidence for the qualitative question of how cues are used by a navigator. For our study, we want to model the use of PI and landmark cues, to gain insights into how the two sources of spatial information interact in the different environmental conditions. We compose our model of a **distance** and a **direction** estimate that can be thought of as the norm and angle of a vector, which represents a complete **position** estimate.

For landmark information, inspired by Jetzschke et al. (2017), we model distance information as Gaussian probability density functions fitted on the distance estimates made by participants relative to the landmark cues (see section “Maximum Likelihood Estimation modelling” for more details):

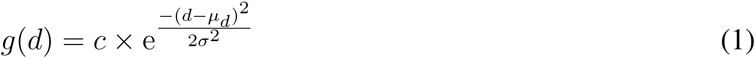

with *d* being the metric distance measure with regard to the landmark, *σ* being the standard deviation of *d*, *µ_d_* its median and 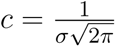 being the scaling factor. Since the PI system not only provides distance but also directional information, we expanded the model to include such a directional component in the form of a circular normal (von Mises) probability density function:

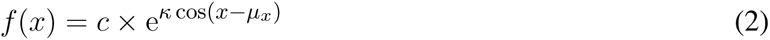

with *x* being angular estimates in the interval [*−π, π*], *µ_x_* being the median of the angular estimates, 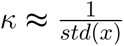 being the concentration parameter and 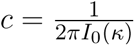 being the scaling factor, where *I*_0_(*κ*) is the modified Bessel function of order zero (see also Murray and Morgenstern (2010) and Zhao and Warren (2015b) for an introduction to modelling directional information using circular functions). Here, the “noisiness” of the cue (variance in the Gaussian) could originate from either perception and the internal spatial representation in encoding the outbound path, or from motor control in executing the homing path. We make no claims towards one source of noise above the others, but consider the PI system as a whole as noisy (Chrastil and Warren, 2017; Harootonian et al., 2020; Chrastil and Warren, 2021; Kessler et al., 2022, preprint). Using these functions, we could derive maximum likelihood estimates for Gaussian (distance) functions centred on every object location, as well as the start of the return trip, which could be combined with a von Mises (direction) function also fitted on the start of the return. This approach to encoding directional estimates has been introduced by Murray and Morgenstern (2010) and employed successfully to model human navigation behaviour by Zhao and Warren (2015b).

Combinations of distance estimates can be calculated by either multiplying or summing the probabilities derived by sampling the functions for each of the landmarks. Here, a multiplication produces an **integration** model 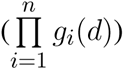, which assumes each cue to be used in a given trial weighted according to its reliability (Zhao and Warren, 2015b; Chen et al., 2017), while a summation produces a cue **alternation** model 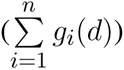, which assumes individual objects are picked at random from trial to trial with selection probabilities based on their reliability (Nardini et al., 2008; Goeke et al., 2016; Chen et al., 2017). For our study, the reliability of all landmarks was assumed to be equal, with the weight derived from the observed reliability in the one object condition (Jetzschke et al., 2017).

For the full model, these combined distance estimates were then always combined with a single direction estimate (*f* (*x*)) centred on the start of the return. This combination was always of the integration form (*L_combined_* = *f* (*x*) *× g_combined_*(*d*)) because a model assuming alternation between using only distance or direction information during a given trial is not plausible given the observed behaviour. To determine goodness of fit, we tested different model variants against the observed data. Specifically, we always determined the MLE for both distance and direction models for the zero object condition, which gave us a model predicting where participants should walk if they followed their PI exclusively. This model was then also evaluated on the data of the other conditions and compared with the variant of the same model fitted on the respective condition. These models were then combined with landmark-based distance models as described above. Parameters for the landmark model were always set to reflect the geometric ground truth of the distances of the three central objects around the goal, and the spread of estimates observed for the one object condition specifically (*µ_d_* = 10, *σ* = *σ̃*(*d*_1_), with *d*_1_ being the distance estimates observed for the first landmark). Model quality was evaluated by calculating the log-likelihood of each model variant (*l* = Σ log(*L_combined_*)). Since model variants were always of the same class, i.e. being composed of the same number and type of functions, model log-likelihoods can be compared directly. Furthermore, we used a bootstrapping procedure to derive confidence intervals for all MLEs, which allows for direct visual comparison of parameters across conditions (see Fig. 9). Finally, we also calculated Akaike and Bayesian information criteria (AIC, BIC) for all models as additional measures for comparison (see supplement “S4 MLE model summaries”). Model rankings remained unchanged regardless of the method of comparison.

#### 2.4.4 Power analysis

For the design of this study, we analysed data from a previous pilot study with a virtual triangle completion task and different numbers of trees (Müller et al., 2023) and did power analyses for repetitions per condition and for number of participants using the one-sample t-test function in G*Power 3.1 (Faul et al., 2007). We planned to collect data from N=20 participants with n=24 repetitions per condition each. Thus, assuming that planned sample size, a power of *β* = 95%, and significance level *α* = 5%, the 2.78 *vm* median standard deviation of position errors within participants and within conditions observed in the pilot study allows to detect differences in position of *≥*2.15 *vm* in a two-sided one-sample t-test. Likewise, the 4.02 *vm* standard deviation of participants’ median position error observed in the pilot within conditions, we can identify differences in position of *≥*3.5 *vm*. This resolution was deemed adequate to resolve the performance differences for our main study.

Using the study’s actual results (N=23, n=24) and following the same calculations, we can actually resolve with the same power, significance level, and t-test, differences of *≥*3.2 *vm* within participants and within conditions, and *≥*2.25 *vm* between participants and within conditions, thus matching and even exceeding the spatial resolution desired for our analysis.

#### 2.4.5 Exclusion criteria

The experiment consisted of training and five main test sessions with initially 25 participants. Six of the participants did not complete the last main session, and thus we decided to include only four main sessions to acquire a balanced dataset and include those six participants that otherwise would have been rejected. Two additional participants were excluded from all analyses, one of which did not finish the experiment in the given time, and one who indicated in the follow-up questionnaire that they were using the computer display’s rim as an external cue to better estimate the goal location. We remained with 23 participants completing 24 trials per condition within four main sessions. From the raw data, we excluded trials with missing values (e.g. due to expired timeout or mistaken indications of the home position due to unintentional double clicks in the in-game UI), which was the case in less than seven trials for any participant. For validation purposes, we also ran our analysis including all available sessions for each participant which yields the same qualitative results with only minor deviations in descriptive statistics. Data processing can be retraced in detail with the supplementary material.

## 3 RESULTS

Healthy participants (N=23) completed a triangle completion homing task in virtual reality. Previous research has shown that the participants’ direction and distance estimates in triangle completion tasks are dependent on triangle size and shape (Loomis et al., 1993; Wiener and Mallot, 2006; Harootonian et al., 2020). Additionally, people showed persistent individual biases in walked angles (Jetzschke et al., 2016). Both of these effects constitute confounding factors in an experiment whose main outcome measure are the distance and angle estimates in question. Therefore, since the focus of this study is the influence of environmental complexity, rather than the nature of these individually different responses to left and right turns and different path geometries, we chose to keep the start point and outbound path constant across all trials and environmental conditions (Fig. 1), forgoing the usual approach of providing mirrored versions of the task with left and right turns. To avoid long-tern learning effects, no feedback as to their performance was provided to participants in test conditions at any time. Nevertheless, performance changes along experimental sessions is discussed where applicable (see section “Disentangling the effect of different degrees of clutter”). While the path shape was kept constant, the number of identical tree objects in the surround was systematically varied between zero and 99 (zero, one, two, three, ten, 99) from trial to trial. Conditions with higher object number always contained the objects of conditions with lower object number as well (Fig. 1). Spatial constellations were kept constant within each condition. This means that the participants always experienced the same constellation of - for example - 99 trees. This is was done to avoid inducing a feeling of low environmental stability in participants, which would have lowered trust in landmark-based guidance cues (Roy et al., 2023; Zhao and Warren, 2015a). To create optic flow for visual estimation of self-motion we provided identical grass tufts on the ground that were randomly arranged for each trial. We quantify homing performance as position error, which is the Euclidean distance between the actual goal location and the endpoint of the trajectory performed by the participant.

### 3.1 General effect of different degrees of clutter

The main measure of interest for our study was the endpoint of each trial’s homing trajectory. This is the location at which the participant estimated the goal location to be. We call the distance from this location to the actual goal location the position error of the trial in question and measure the distance in virtual meters (VM). Virtual meters resemble actual meters in the VR, based on the scaling of visible 3D assets (grass tufts, trees, wood planks, campfire).

We find that the number of objects (trees) present in a given trial substantially impacts the position error (Fig. 2). Both median position error (homing accuracy), as well as the spread in errors (homing precision) differ greatly for different environments. Specifically, when no objects are available, we observe a large median error of almost 10 virtual meters (*vm*), *±* 6 *vm* (standard deviation) (see Fig. 3 and 4). When individual objects are added to the scene, homing accuracy increases, as the median position error decreases to a minimum of little more than 2 *vm* for trials in which the goal location was constrained by three surrounding objects. Precision also increases with a median standard deviation of position error as small as 2 *vm*. However, if further objects are added beyond the initial three, a noticeable trend emerges: for numerous participants, the position error once again rises as the object number reaches ten or 99 (e.g. Fig. 2b). This trend is accompanied by a considerable increase in the standard deviation of position errors among different participants (as indicated by the marker colour in Fig. 2 and 4). Consequently, this leads to an overall decrease in precision, bringing it to a level similar to what is observed for environments devoid of any objects. Nevertheless, for several participants, the accuracy and precision stay roughly at the level observed for the three object condition (e.g Fig. 2a), with one participant even showing the highest precision in the 99 object condition (Fig. 4, bottom). Hence, the number of objects influences performance of individual participants differently. We observe this strong effect of object number on accuracy and precision, although participants walked the same outbound path in all trials. This highlights the strength of the effect of object number, as the consistency of the outbound path should only be able to attenuate differences between different object numbers, rather then increase them. While we describe the patterns of errors in ascending order from few to many objects present in the visual scene, conditions were presented in random order to the participants during the experiment.

**Figure 2.**
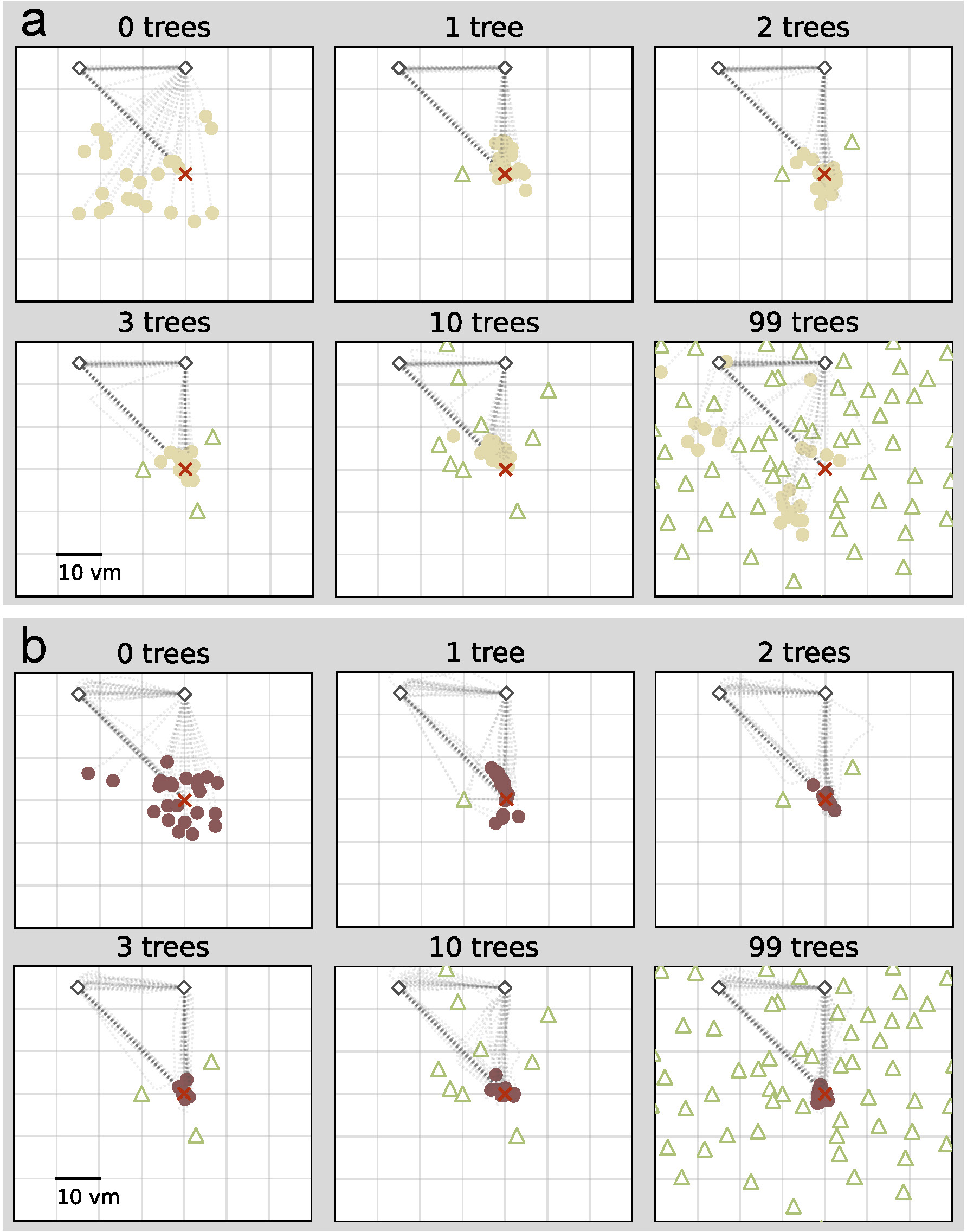
Homing performance of two example participants depends on number of objects. For both participants performance increases with up to three objects. For one participant (a), performance decreases with ten or 99 objects, and for another it stays high (b). Trajectory endpoints of single trials are depicted by dots. Colours marking individual Participants are matched across all figures, based on a given participant’s median position error in the 99-object condition (refer to Fig. 4 top right). Outbound and homing trajectories are indicated by light grey dotted lines. Marker size does not match actual tree dimensions. Cutout does not show all objects, please see Fig. 1 for a more zoomed-out view of the environment.

**Figure 3.**
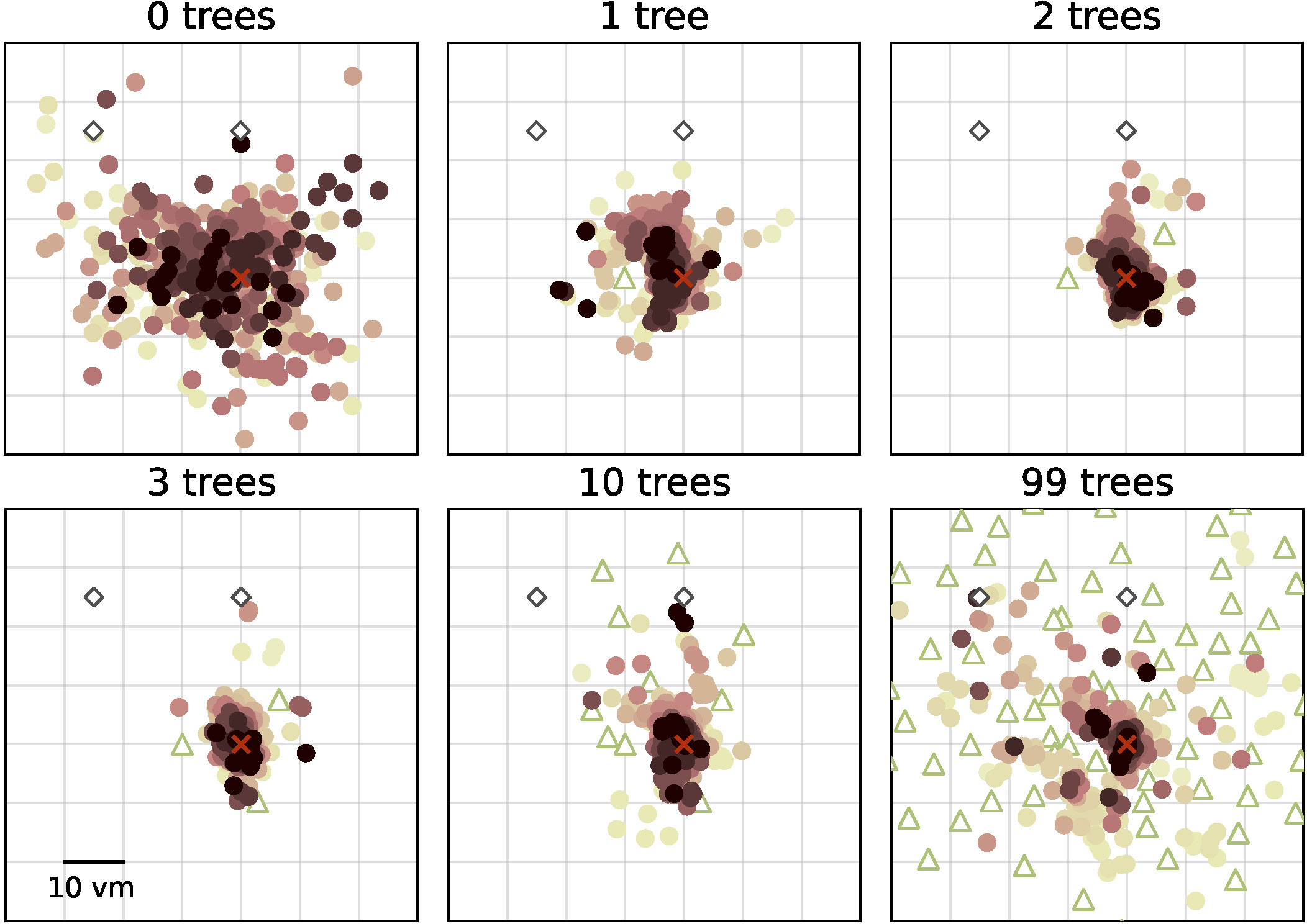
Homing performance at population level increases in environments with up to three objects, and then decreases up to 99 objects. Trajectory endpoints of single trials are depicted by dots. Colours marking individual Participants are matched across all figures, based on a given participant’s median position error in the 99-object condition (refer to Fig. 4 top right). Marker size does not match actual tree dimensions. Cutout does not show all objects, please see Fig. 1 for a more zoomed-out view of the environment.

**Figure 4.**
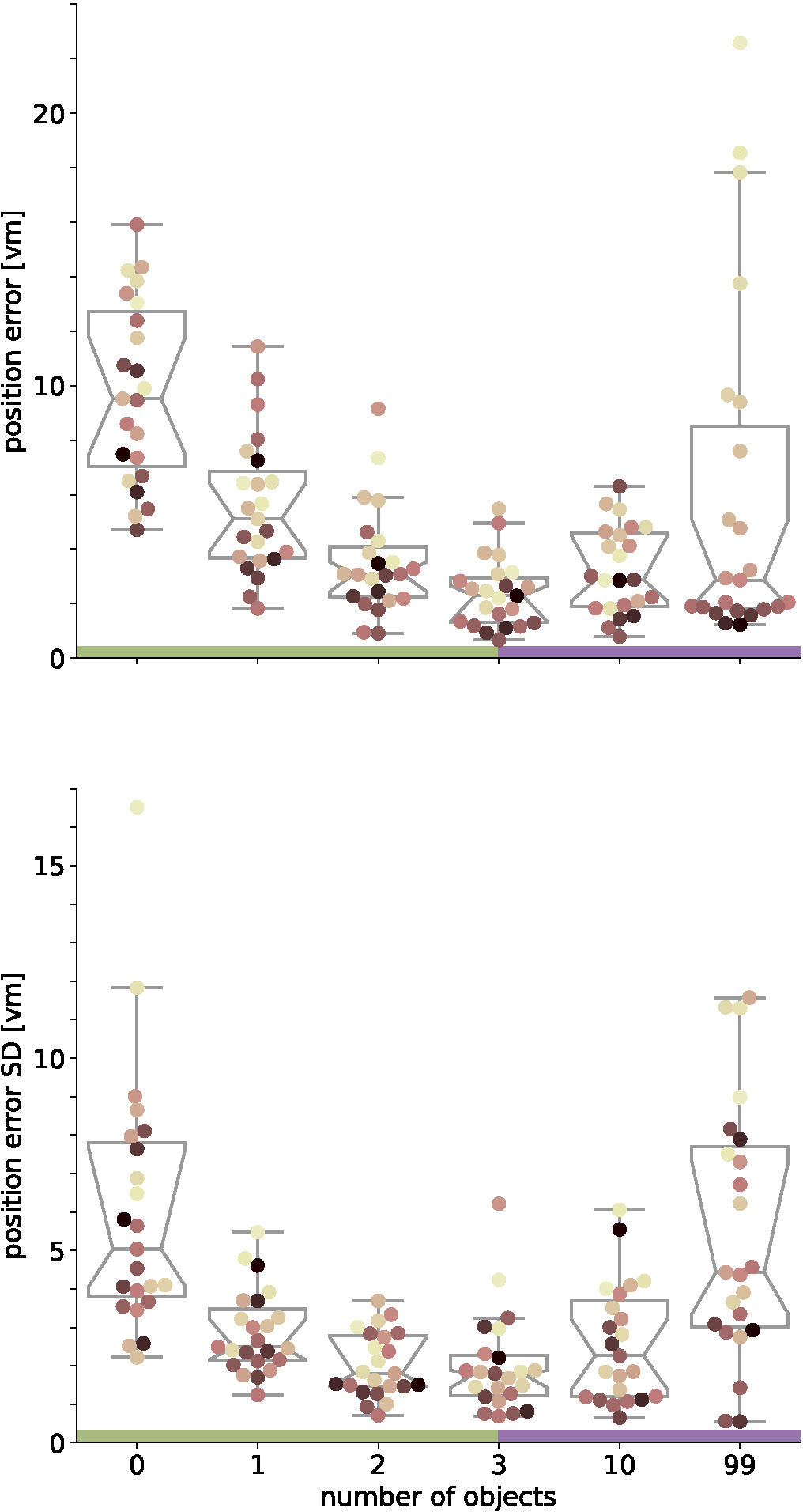
Position error accuracy (top) and precision (bottom) on population level are lowest with three objects. Boxplots are based on median position error in virtual meters (*vm*) (top) or standard deviation (SD) (bottom) of N=23 single participants with n=24 repetitions within respective conditions (see dots). Colours marking individual Participants are matched across all figures, based on a given participant’s median position error in the 99-object condition (refer to Fig. 4 top right). Boxplot notches indicate 95% confidence intervals around the median. Green and purple horizontal bars at the bottom indicate low and high clutter effects (see section “Separating effects of low and high clutter” for details)

### 3.2 Disentangling the effect of different degrees of clutter

We fit an initial linear mixed effects (LME) model using only the experimental condition (number of objects) as an ordinal predictor, which already showed significant predictive power for both accuracy (Tab. 1 model 1) and precision (Tab. 1 model 4) of goal estimates (see Methods “Linear mixed effects models” for methodological details). To investigate possible individual differences in performance across conditions, we added participant identity as a random predictor (Tab. 1 models 2 and 4), which significantly improved the model for accuracy, and for precision (likelihood ratio test: accuracy: *χ*^2^ ≈21.85, logLik1=-112.5, logLik2=-122.4, p*<*0.0001; precision: *χ*^2^ ≈20.49, logLik1=-202, logLik2=-212.2, p*<*0.0001).

**Table 1.**
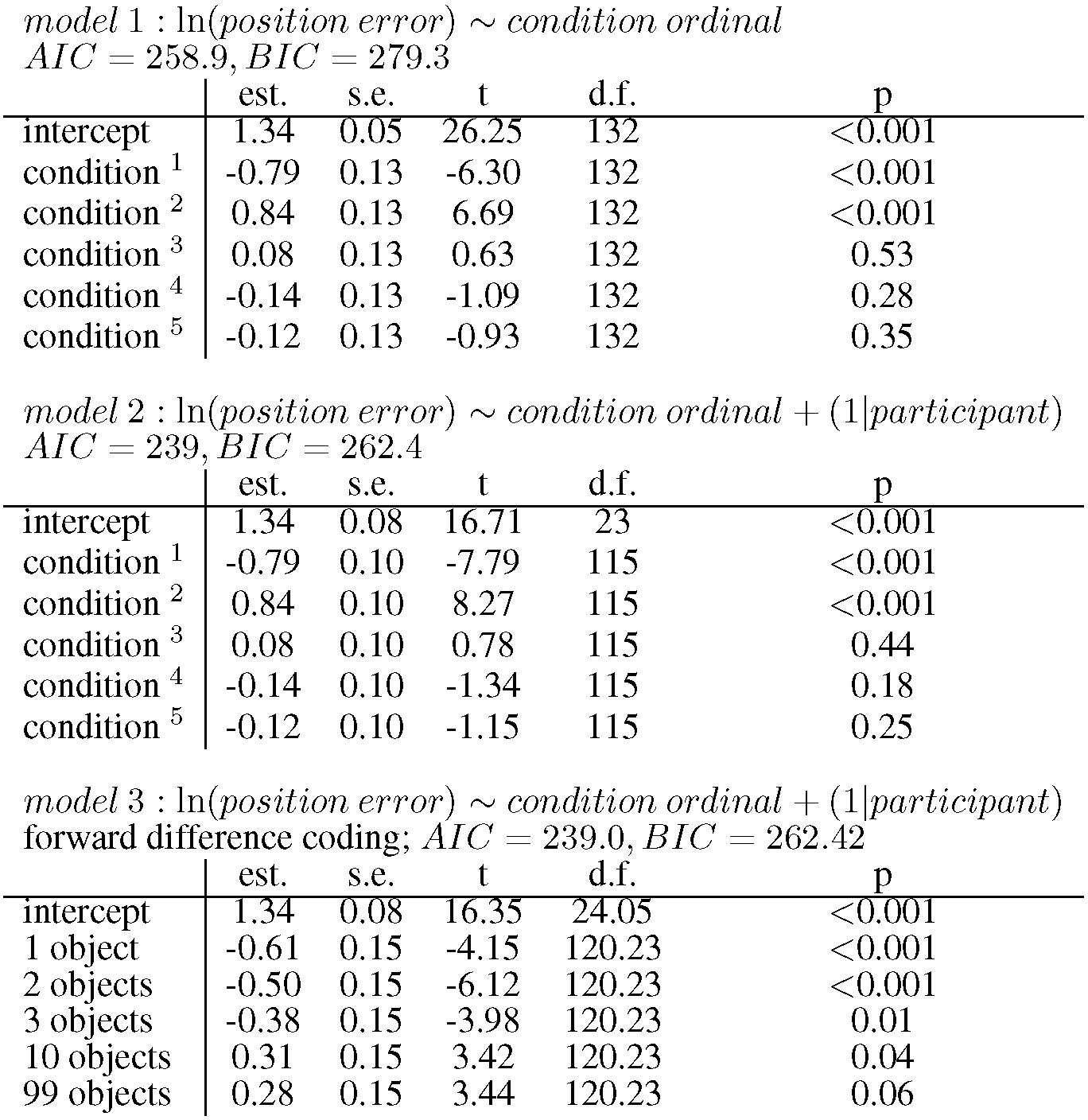
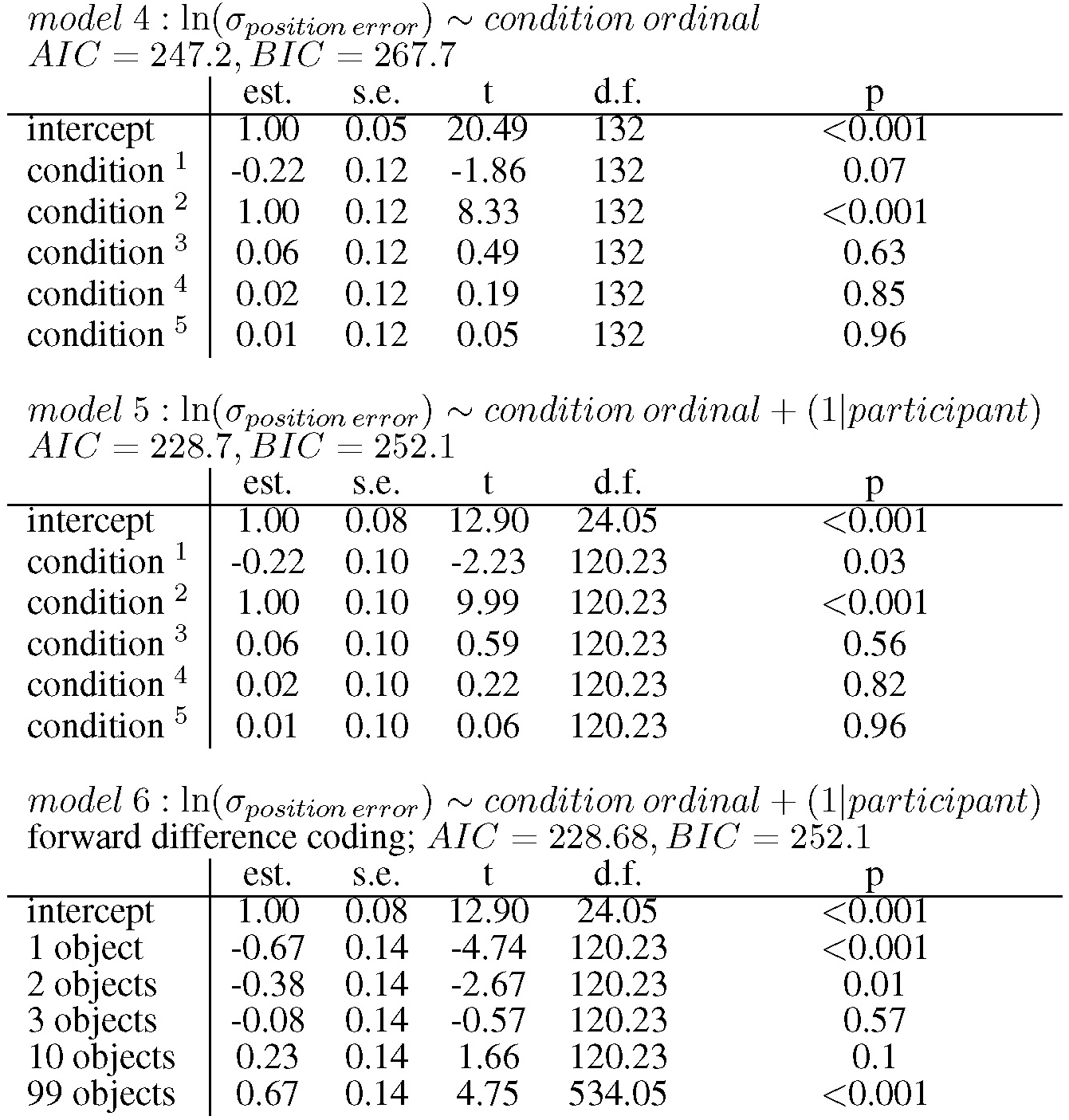
Performance of linear regression models with and without the random effect of participant. Model 1 predicts the position error from the condition (number of objects). Model 2 additionally includes a random participant factor. Model 3 restates model 2 but uses forward difference coding. Models 4 to 6 follow the same structure as 1 to 3, but predict position error variance instead. In all models condition is ordinal and polynomials from 1st to 5th degree are fitted. est.: estimate; s.e.: standard error; t: t value; d.f.:degrees of freedom; p: p-value

|  | est. | s.e. | t | d.f. | p |
| --- | --- | --- | --- | --- | --- |
| intercept | 1.34 | 0.05 | 26.25 | 132 | <0.001 |
| condition <sup>1</sup> | -0.79 | 0.13 | -6.30 | 132 | <0.001 |
| condition <sup>2</sup> | 0.84 | 0.13 | 6.69 | 132 | <0.001 |
| condition <sup>3</sup> | 0.08 | 0.13 | 0.63 | 132 | 0.53 |
| condition <sup>4</sup> | -0.14 | 0.13 | -1.09 | 132 | 0.28 |
| condition <sup>5</sup> | -0.12 | 0.13 | -0.93 | 132 | 0.35 |

|  | est. | s.e. | t | d.f. | p |
| --- | --- | --- | --- | --- | --- |
| intercept | 1.34 | 0.08 | 16.71 | 23 | <0.001 |
| condition <sup>1</sup> | -0.79 | 0.10 | -7.79 | 115 | <0.001 |
| condition <sup>2</sup> | 0.84 | 0.10 | 8.27 | 115 | <0.001 |
| condition <sup>3</sup> | 0.08 | 0.10 | 0.78 | 115 | 0.44 |
| condition <sup>4</sup> | -0.14 | 0.10 | -1.34 | 115 | 0.18 |
| condition <sup>5</sup> | -0.12 | 0.10 | -1.15 | 115 | 0.25 |

*model 3 : $\ln(\text{position error}) \sim \text{condition ordinal} + (1|\text{participant})$* *forward difference coding; $AIC = 239.0, BIC = 262.42$*
|  | est. | s.e. | t | d.f. | p |
| --- | --- | --- | --- | --- | --- |
| intercept | 1.34 | 0.08 | 16.35 | 24.05 | <0.001 |
| 1 object | -0.61 | 0.15 | -4.15 | 120.23 | <0.001 |
| 2 objects | -0.50 | 0.15 | -6.12 | 120.23 | <0.001 |
| 3 objects | -0.38 | 0.15 | -3.98 | 120.23 | 0.01 |
| 10 objects | 0.31 | 0.15 | 3.42 | 120.23 | 0.04 |
| 99 objects | 0.28 | 0.15 | 3.44 | 120.23 | 0.06 |

Table 1. continued. *model 4 : $\ln(\sigma_{\text{position error}}) \sim \text{condition ordinal}$ $AIC = 247.2, BIC = 267.7$*
|  | est. | s.e. | t | d.f. | p |
| --- | --- | --- | --- | --- | --- |
| intercept | 1.00 | 0.05 | 20.49 | 132 | <0.001 |
| condition <sup>1</sup> | -0.22 | 0.12 | -1.86 | 132 | 0.07 |
| condition <sup>2</sup> | 1.00 | 0.12 | 8.33 | 132 | <0.001 |
| condition <sup>3</sup> | 0.06 | 0.12 | 0.49 | 132 | 0.63 |
| condition <sup>4</sup> | 0.02 | 0.12 | 0.19 | 132 | 0.85 |
| condition <sup>5</sup> | 0.01 | 0.12 | 0.05 | 132 | 0.96 |

*model 5 : $\ln(\sigma_{\text{position error}}) \sim \text{condition ordinal} + (1|\text{participant})$ $AIC = 228.7, BIC = 252.1$*
|  | est. | s.e. | t | d.f. | p |
| --- | --- | --- | --- | --- | --- |
| intercept | 1.00 | 0.08 | 12.90 | 24.05 | <0.001 |
| condition <sup>1</sup> | -0.22 | 0.10 | -2.23 | 120.23 | 0.03 |
| condition <sup>2</sup> | 1.00 | 0.10 | 9.99 | 120.23 | <0.001 |
| condition <sup>3</sup> | 0.06 | 0.10 | 0.59 | 120.23 | 0.56 |
| condition <sup>4</sup> | 0.02 | 0.10 | 0.22 | 120.23 | 0.82 |
| condition <sup>5</sup> | 0.01 | 0.10 | 0.06 | 120.23 | 0.96 |

*model 6 : $\ln(\sigma_{\text{position error}}) \sim \text{condition ordinal} + (1|\text{participant})$ forward difference coding; $AIC = 228.68, BIC = 252.1$*
|  | est. | s.e. | t | d.f. | p |
| --- | --- | --- | --- | --- | --- |
| intercept | 1.00 | 0.08 | 12.90 | 24.05 | <0.001 |
| 1 object | -0.67 | 0.14 | -4.74 | 120.23 | <0.001 |
| 2 objects | -0.38 | 0.14 | -2.67 | 120.23 | 0.01 |
| 3 objects | -0.08 | 0.14 | -0.57 | 120.23 | 0.57 |
| 10 objects | 0.23 | 0.14 | 1.66 | 120.23 | 0.1 |
| 99 objects | 0.67 | 0.14 | 4.75 | 534.05 | <0.001 |

The residuals in error scores for both models are characterised by a degree of heteroscedasticity, which is caused mainly by the large variance in the 99 object condition (see Fig. 4). Accordingly, residuals for errors are homoscedastic across conditions when the 99 object condition is excluded. However, as moderate heteroscedasticity is not considered to be a major influence on LME models (Schielzeth et al., 2020), we consider the model family described above as adequate to describe the observed data. Thus, we base our analysis of accuracy on a regression model of the following form:

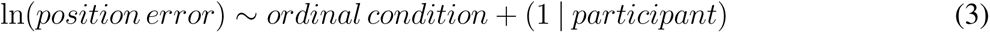

and for the analysis of precision in this form:

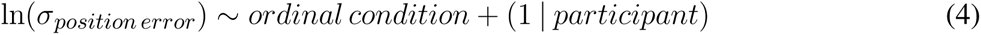

with *σ* being the observed standard deviation of the position error. While the model explains approximately *R*^2^ ≈ 59% of variance in our median position errors, 40% are attributed to our fixed condition effect. For position error variance approximately *R*^2^ ≈ 55% of variance is explained, 36% are attributed to the condition effect. Note that there is no way to calculate *R*^2^-values for mixed effects models, and the values reported above are approximations based on an implementation of Nakagawa and Schielzeth (2013) with refinements by Johnson (2014).

The model suggests a linear as well as a quadratic component in the relationship between condition and position error for accuracy (Tab. 1 model 2) and precision (Tab. 1 model 5) that we investigate in more detail below.

We observe a trend of reduced position error over sessions (see supplement “S2 Session order effect”), however, including session number as an additional fixed effect in the regression model results in only a slight improvement of fit (without session: AIC≈1163.4, BIC≈1193.6; with session: AIC≈1150.9, BIC≈1194; likelihood ratio test: *χ*^2^ ≈18.58, logLik1=-574.7, logLik2=565.4, p≈0.00033). Since the learning effect is small compared to the effect of object number (see supplement “S2 Session order effect”), and learning is not the primary focus of this study, we do not consider the effect in further analyses.

### 3.3 Separating effects of low and high clutter

We set out to investigate the effects of environmental complexity on navigation performance. In our experiments, complexity was manipulated by changing the number of objects present in the virtual environment. Based on the patterns of performance described above, we will group our experimental conditions into low clutter effects for trials with zero to three objects (green bar in Fig. 4 and 5, and green distributions in Fig. 6), and high clutter effects for trials with three to 99 objects (purple bar in Fig. 4 and 5, and purple distributions in Fig. 6) respectively. We motivate this separation firstly based on a local minimum in a quadratic regression at the three object condition (see Fig. 5), and secondly based on the change of sign at the three object condition in a forward difference contrast coding (UCLA Statistical Consulting Group, 2021) version of our regression model (see Tab. 1 and supplement “S3 Separating effects of low and high clutter” for details).

**Figure 5.**
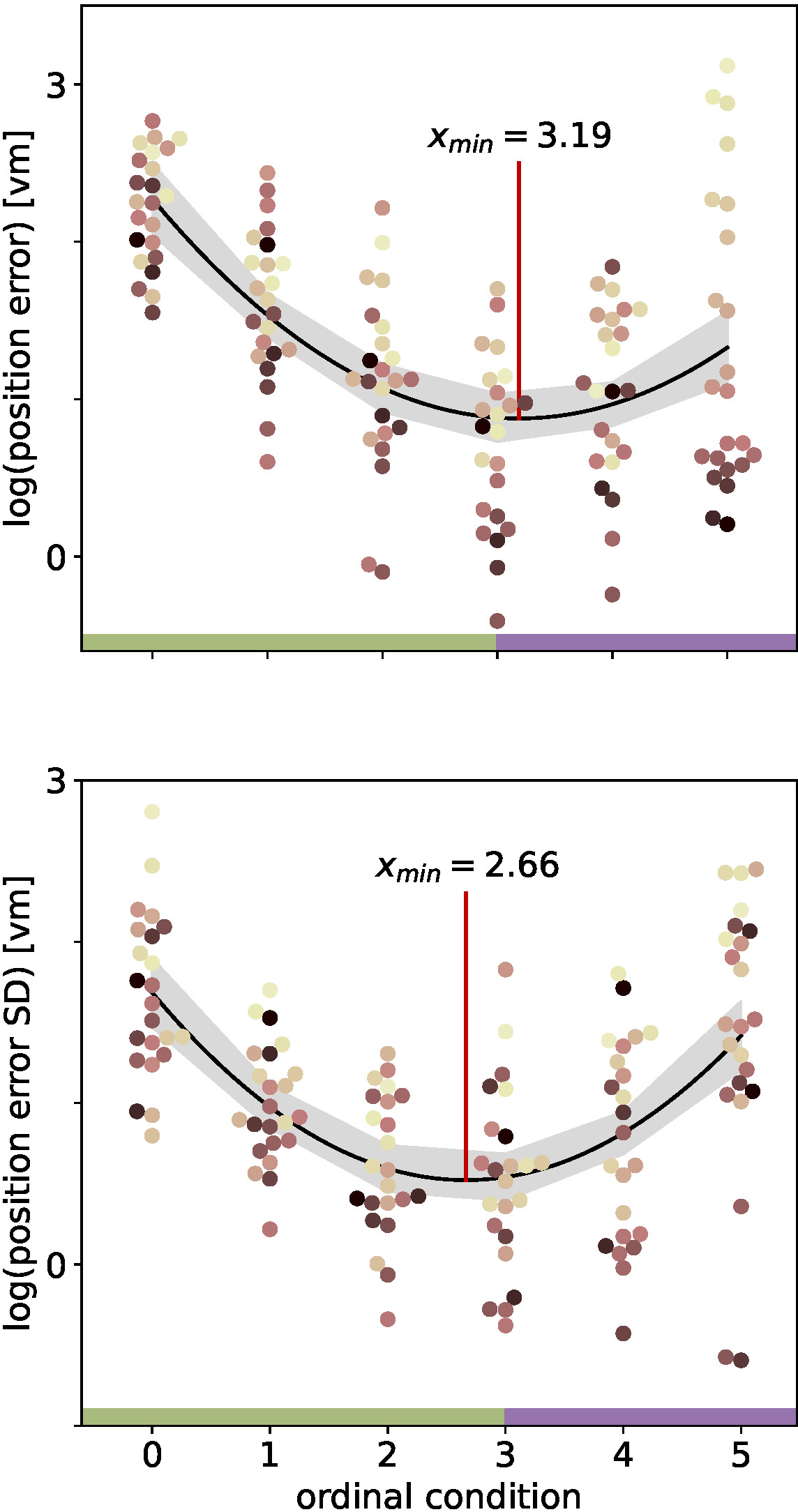
Quadratic fits on the log-transformed position error (top) or log-transformed position error standard deviation (SD) (bottom) with ordinal condition show a local minimum (red vertical line) close to the three object condition. Fits include data from N=23 participants with n=24 repetitions per condition. Ordinal conditions from zero to five correspond to conditions with zero, one, two, three, ten, and 99 objects respectively. Quadratic regressions fit the data significantly better than a linear regression and the local minimum at three objects is validated in a linear regression model with forward difference coding (see Tab. 1). Colours marking individual Participants are matched across all figures, based on a given participant’s median position error in the 99-object condition (refer to Fig. 4 top right). Green and purple horizontal bars at the bottom indicate low and high clutter effects (see section “Separating effects of low and high clutter” for details)

**Figure 6.**
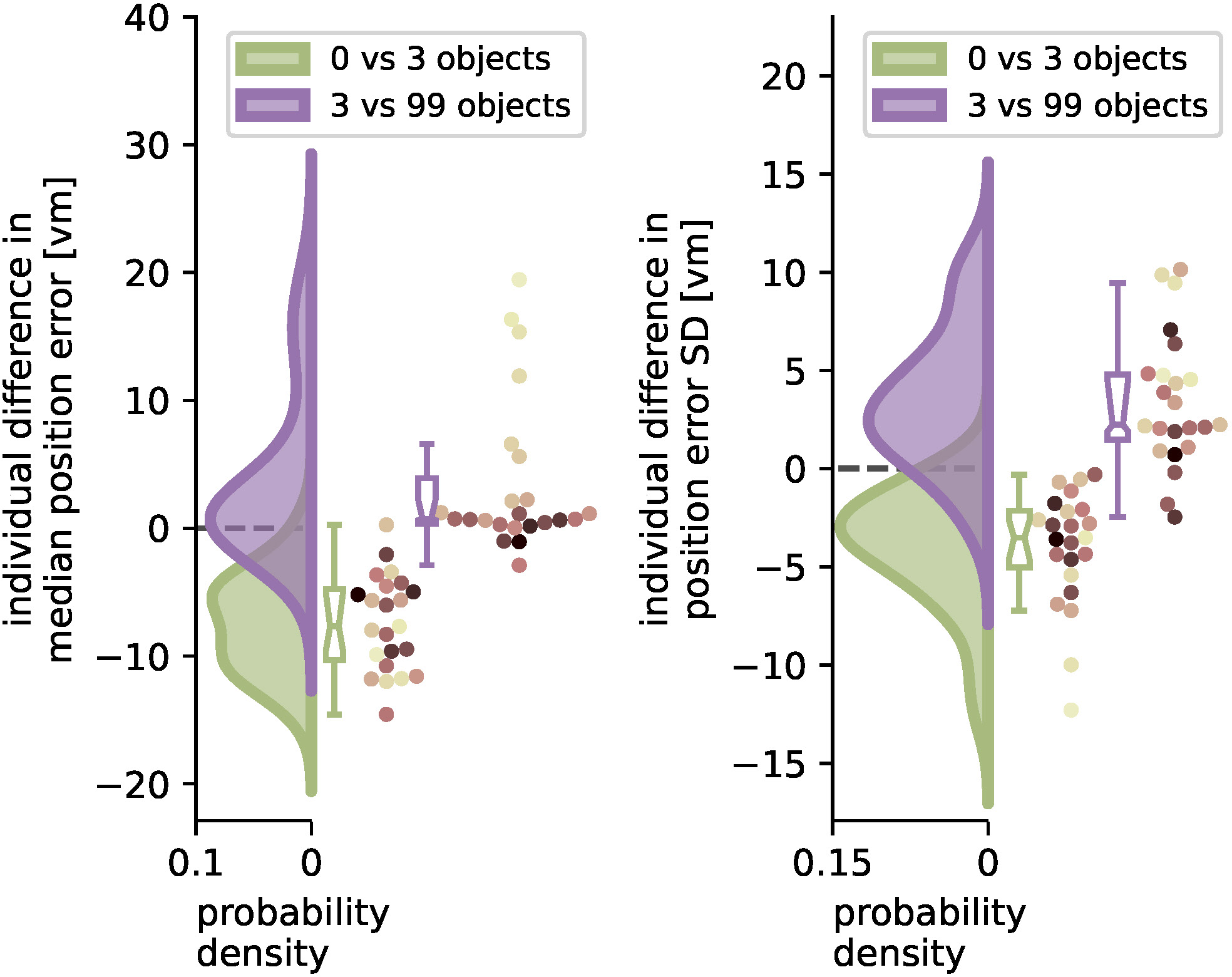
Performance improvement in accuracy and precision of position error from zero to three objects and performance impairment from three to 99 objects. Green and purple curves indicate kernel-density estimations fitted to the difference in median position error (left) or position error standard deviation (SD) (right) between zero and three objects (green), or three and 99 objects (purple), respectively, of N=23 single participants with n=24 repetitions within each condition (see dots). Boxplot notches indicate 95% confidence intervals around the median. All but one participant performed better with three than with zero objects. 11 participants performed significantly worse, and 3 significantly better with 99 objects, while 10 others remained equally good (see Results “Quantifying effects of low and high clutter” for details). Colours marking individual Participants are matched across all figures, based on a given participant’s median position error in the 99-object condition (refer to Fig. 4 top right).

### 3.4 Quantifying effects of low and high clutter

With the three object condition separating low and high clutter effects, we ask which of the factors (number of objects and individual differences) can better explain the changes in homing performance for more and less cluttered environments. To quantify these effects, we analyse the median performance change in accuracy and precision from the zero object to the three object condition and compare it to the change from the three object to the 99 object condition for each individual participant. Note that this analysis was conducted without log-transforming position error because it is not subject to assumptions of linear regression models, and offers easier interpretability of the results.

When comparing the two distributions of performance changes in the low clutter and high clutter environments (see Fig. 6) we see that, as expected, the position error and its standard deviation decrease as objects are added in the “low clutter” (zero to three objects) environments. Conversely, in “high clutter” (three to 99 objects) environments, we find in the kernel density distribution for accuracy one local maximum at 0 *vm* and a second lower one around 15 *vm* (Fig. 6 left), and for precision a local maximum at 2 *vm* and a heavy tail going beyond 10 *vm*, respectively (Fig. 6 right). The first peak indicates no considerable change in position error for most participants, while the second one depicts participants, for whom 99 objects lead to an impairment of homing performance. To capture this group we chose the bootstrapped 95% confidence interval of the median of the distribution of differences. Seven out of 23 participants, show a performance decline in accuracy between three and 99 objects outside the bootstrapped 95% confidence interval and thus fall into this group of people negatively impacted by the high degree of spatial clutter. Conversely, three other participants even perform slightly better with 99 objects than with three. This attribution was also confirmed using a Wilcoxon ranksum test on single trial basis for significant changes in performance between three and 99 objects (11 get sign. worse (all p*<*0.04, −2.88*<*d*<*-0.65), 3 get sign. better (all p*<*0.016, 0.05*<*d*<*0.23), 10 being insignificant). Correspondingly, the distribution of median accuracy differences in position error between three and 99 objects differs significantly from normality (Shapiro-Wilk test: W≈0.735, p*<*0.0001), while the distribution of differences between zero and three objects does not (Shapiro-Wilk test: W≈0.974, p≈0.793).

We therefore conclude that the number of objects is the main driver of participant homing performance in low clutter conditions (zero to three objects), while for the high clutter conditions (three to 99 objects) the individual behaviour of the participants plays a larger role in determining their success.

### 3.5 Relation between accuracy and precision

We have analysed median homing errors (accuracy) and the standard deviation of errors (precision) and found matching patterns for environments with different degrees of clutter for both measures. This is supplemented by direct correlations of the two measures in all conditions (see Fig. 7), which shows that higher accuracy is usually associated with higher precision, at least on the population level. However, looking at the performance of single participants reveals inter-individual differences. Participants exhibit different degrees of accuracy and precision, with some being accurate but imprecise (Fig. 7a and e), and others being inaccurate but precise (Fig. 7b and c). These participants weaken the precision-accuracy correlation at the population level. The strongest correlation was found in the zero object condition, where 44% of variance in accuracy is explained by precision.

**Figure 7.**
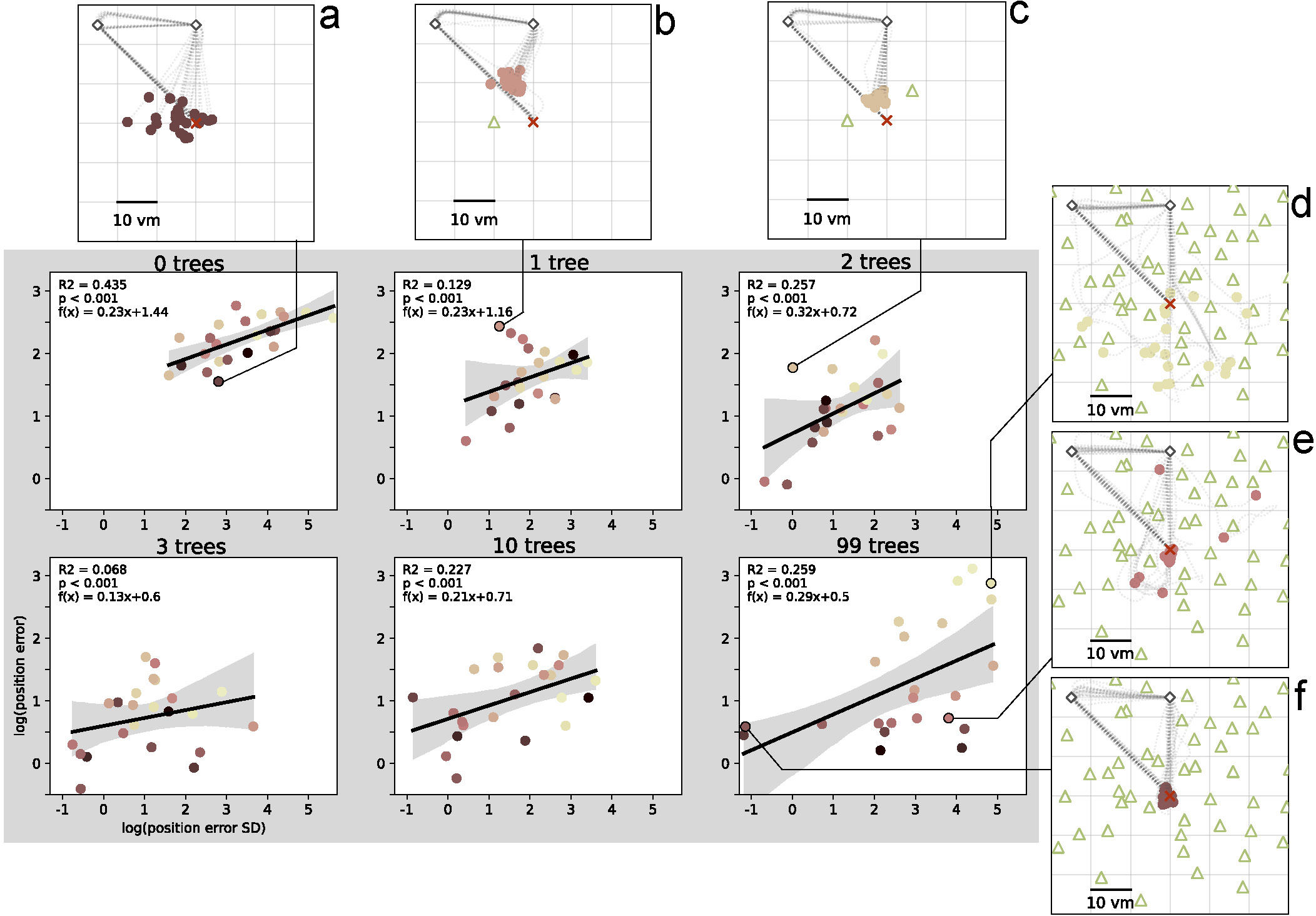
Position error precision and accuracy are significantly correlated in all conditions. There are large inter-individual differences with example participants (a-f) respectively performing accurately and imprecisely (a and e), inaccurately and precisely (b and c), inaccurately and imprecisely (d), or accurately and precisely (f). Data points indicate median position error and its standard deviation (SD) for each of N=23 participants with n=24 repetitions per condition. Colours marking individual Participants are matched across all figures, based on a given participant’s median position error in the 99-object condition (refer to Fig. 4 top right). Grey areas indicate 95% confidence intervals. Marker size does not match actual tree dimensions.

### 3.6 Modelling cue interactions using MLE

To capture the interplay between landmark guidance and PI in our study, we have developed an MLE model that builds on the models of Jetzschke et al. (2017) for distance estimates and Zhao and Warren (2015b) for directional estimates, who each found a good fit for their models with behavioural data for up to three landmarks. Our model combines the distance estimation from landmark cues, proposed by Jetzschke et al. (2017), with model components representing additional distance estimation (see Fig. 8A) and direction estimation (see Fig. 8B) from PI cues (see methods “Maximum Likelihood Estimation modelling” for details). The encoding of directional estimates is based on Zhao and Warren (2015b). In addition to combining these previous partial models to fully encode the spatial information available to our participants, we challenge the resulting combination model in conditions with ten and 99 objects. By either setting or fitting different values to the parameters of the three model components (landmark-based distance estimate, PI-based distance and direction estimate) and varying between two types of combination (integration or alternation of objects), we can quantitatively express different hypotheses about how the landmarks and PI are used by the participants.

**Figure 8.**
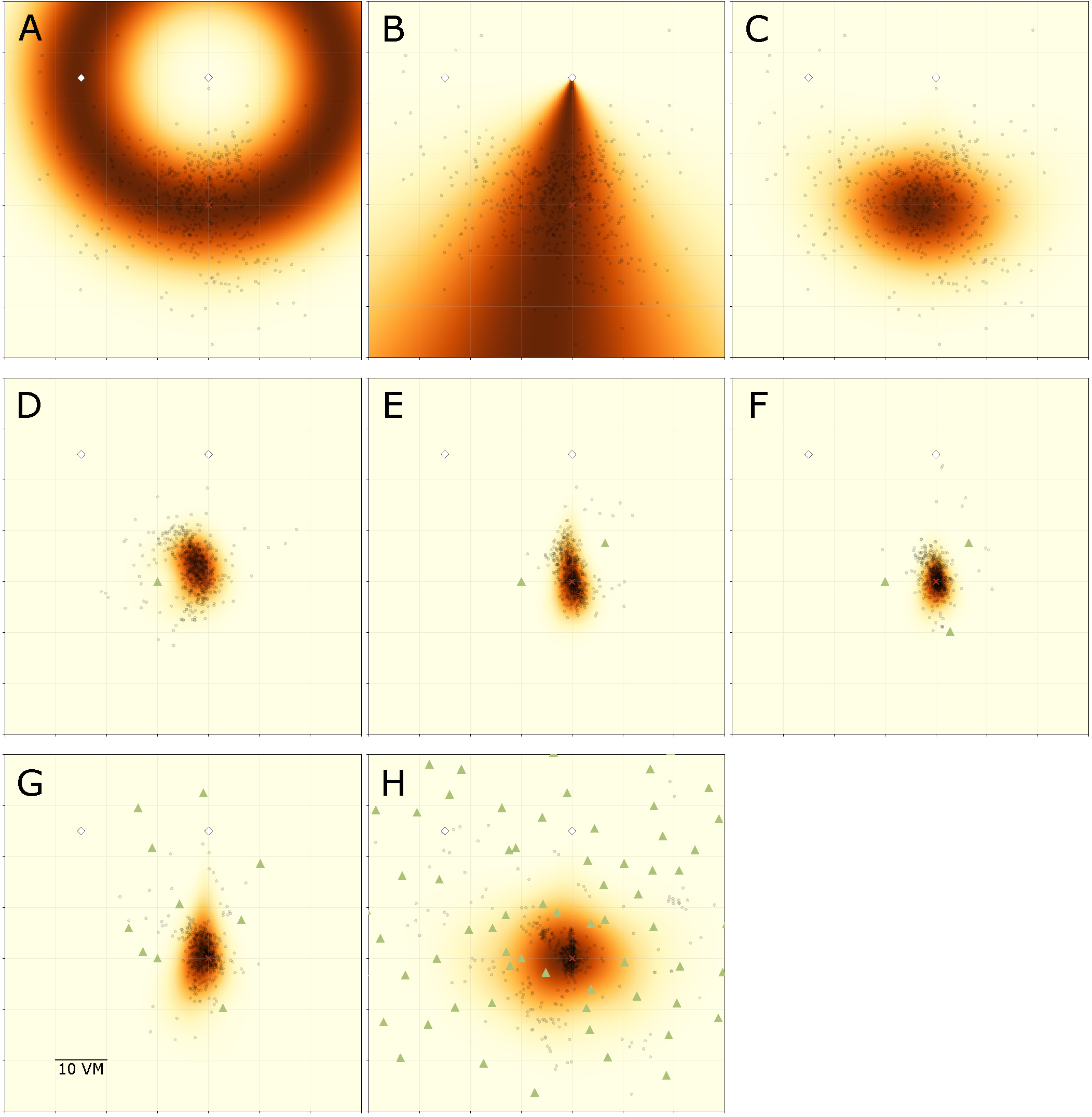
Overview of MLE modelling results. Panels show participant goal estimates (black dots) and landmark positions (green triangles) for different experimental conditions, overlayed on top of maximum likelihood model predictions for the respective condition. Heatmap colour signifies the predicted likelihood of targeting a given area, with darker colours indicating greater likelihood. Marker size does not match actual tree dimensions. Panel A introduces the ring-shaped distance estimate model, based on fitting a normal distribution to the distance component of the goal estimates. Heatmap shows the overall best model fit (maximum likelihood estimate) for this model type for the pooled data of all N=23 participants for the zero object condition. Panel B introduces the fan-shaped direction estimate model, based on fitting a von Mises (circular normal) distribution to the directional component of the goal estimates. Heatmap shows the overall best model fit (maximum likelihood estimate) for this model type for the pooled data of all participants for the zero object condition. Panel C shows the prediction resulting from an integration of both model components (*ring × fan*) for the same data. Panels D-F show the resulting prediction from an integration model combining a full PI model with an additional ring model for each object fitted at 10 *vm* distance (*ring × fan ×* Π *ring*). Panels G-H show the resulting prediction from a model combining a full PI model with an alternation of ring models for each object fitted at 10 *vm* distance (*ring × fan ×* Σ *ring*).

We summarise the model components in Tab. 2 and visualise the spatial combination of functions in Fig. 8, which shows the result of performing MLE along both the distance and the angular domains. It can be clearly seen, how the PI-driven angular estimate provides the spatial constraint needed to limit position estimates from the full ring given by the landmark-driven distance function alone to the area actually targeted by participants.

**Table 2.** Overview of maximum likelihood estimation model components. “*g*” denotes distance models, “*f*” denotes direction models. “*LM*” and “*PI*” denote whether models were based on the landmark information or PI estimate respectively. “*∗*” denotes that models parameters were maximum likelihood estimates, either for the current condition or for the 0-object-condition (0 *objs*) if specified. “*true — dist*” denotes that landmark models were fitted using the true distance to the goal for the 3 central objects surrounding the goal, rather than the MLE for the given data. “Π” denotes integration of individual landmark models, while “Σ” denotes alternation. Top subdivision lists basic model components, middle subdivision gives the form of the full models used, bottom subdivision gives the best performing model variant for the different conditions.

| model | description |
| --- | --- |
| $g_{PI}$ | PI-based distance model component |
| $f_{PI}$ | PI-based direction model component |
| $g_{LM}$ | LM-based distance model component |
| $g_{PI} \times f_{PI} \times \prod g_{LM}$ | integration model example |
| $g_{PI} \times f_{PI} \times \sum g_{LM}$ | alternation model example |
| $g_{PI}^* \times f_{PI}^*$ | best fit 0 objects |
| $g_{PI}^* \times f_{PI}^* \times \prod g_{LM}^{\text{true-dist}}$ | best fit 1-3 objects |
| $g_{PI}^* \times f_{PI}^* \times \sum g_{LM}^{\text{true-dist}}$ | best fit 10-99 objects |

In the zero object condition, there is only one source of distance and direction information, namely the PI system (see Fig. 8 top row). However, each added object provides another source of distance information relative to itself, which we incorporate into our model. Specifically, each of the three objects present in the one to three object conditions had a goal distance of 10 *vm*. These objects surround the goal location and allow for very precise homing in all conditions in which they are present, if identified correctly among the other objects, as we describe above (see section “Introduction”).

Model comparisons show that model variants including the information from PI always outperform those disregarding the PI information when trying to predict participant behaviour (best models for each condition shown in Fig. 8 middle and bottom row, full comparison of model fits in supplement “S4 MLE model summaries”). This clearly indicates that participants combine information provided by their PI system with information provided by the objects to improve their homing performance. It is especially intriguing, that the prediction areas produced by these models very closely match the observed participant behaviour without needing to fit the parameters of the landmark models (see Fig. 8, panels E-G), which are instead based on the function mean set at the geometrically correct distance of 10 *vm*. This kind of match indicates that 1) the assumptions made by the models represent the actual navigation strategies employed by the participants to a reasonable degree and 2) that in the one to three landmark conditions the participants’ goal estimate is, on average, identical to the geometrically correct goal location.

For the high-clutter conditions (ten, 99 objects, Fig. 8, panels H-I), a full integration approach, as described above, is not sensible, neither from a conceptual nor from a mathematical perspective. From a conceptual standpoint, it is unlikely that participants explicitly memorise distances to such large numbers of objects and combine the information across the whole environment. Mathematically, the integration of such a large number of cues, which involves the multiplication of probabilities, results in likelihoods approaching negative infinity, precluding any meaningful comparison operations among variants of models within this class. As an alternative first approach to capturing navigation behaviour in these more cluttered environments, we used a cue alternation approach (see methods “Maximum Likelihood Estimation modelling” for details). This model class is conceptually more appropriate for the ten and 99 object conditions, since we observe a broad spread of responses across the environment in this condition. Such a pattern matches the prediction of cue alternation models, which lack the variance reduction over single cue conditions predicted by integration models (Chen et al., 2017). As implemented, our alternation model variants assumed participants mistook different trees as the “correct” three trees around the goal in different trials. We do not assume tree actual mis-identification to be completely random, but in the absence of direct evidence for the use of any specific trees, a random selection is sensible as a first approach to capturing an alternation between different trees as spatial cues (for a discussion of the implications of this encoding scheme, see Discussion section “The forest or the trees?”).

For our analysis, we have evaluated integration and alternation models for each condition and compared the goodness of fit of variants using different model parameters. Specifically, we compare models fitted on the results of the zero object condition, which represents homing performance driven purely by PI, with those fitted on the data of each of the other conditions (see Tab. 2 for an overview and supplement “S4 MLE model summaries” for detailed comparisons). If the use of PI was unchanged across conditions, these model variants should not differ in goodness of fit. However, we find that the zero object models are always outperformed by the models fitted on the data for the given condition, when evaluated on the data of the given condition. This indicates that the presence of objects does not only add further information in the form of a second type of cue but also improves the quality of the information provided by the PI system, possibly due to stronger optic flow with more visual structure in the scene. This assessment is further supported by the fact that when considering the PI components of our models, parameter estimates qualitatively differ between the landmark and no-landmark conditions, which should not be the case if PI-use remains unchanged across conditions. For the average PI-based distance estimate, we observe a degree of *overshooting* on the population level for the zero object condition, while a degree of *undershooting* is observed for the conditions in which landmarks were present (Fig 9 top left panel). Intriguingly, the overshoot in distance estimates returns for the 99 object condition, matching the results of the analysis of position errors presented above, which indicate that several participants behave very similarly in the zero object and 99 object conditions. We find similar results for the PI-based direction estimates, where on the population level, participants overshoot the homing direction of 90*^◦^*. This result is in qualitatively agreement with previous studies showing that small angles are often overshot and large angles are undershot (Loomis et al., 1993; Chrastil and Warren, 2017) (but note that Harootonian et al. (2020) did not find a regression-to-the-mean effect). Our results indicate that at least two objects seem to be required to overcome the directional bias observed in the no-landmark condition (Fig 9 bottom left panel) for various participants. When investigating the spread of estimates in both distance and direction (Fig 9 right panels), we find that results closely match those observed in the analysis of position errors, with precision increasing (spread in estimates decreasing) as up to three landmarks are added and then decreasing again as further objects are added.

**Figure 9.**
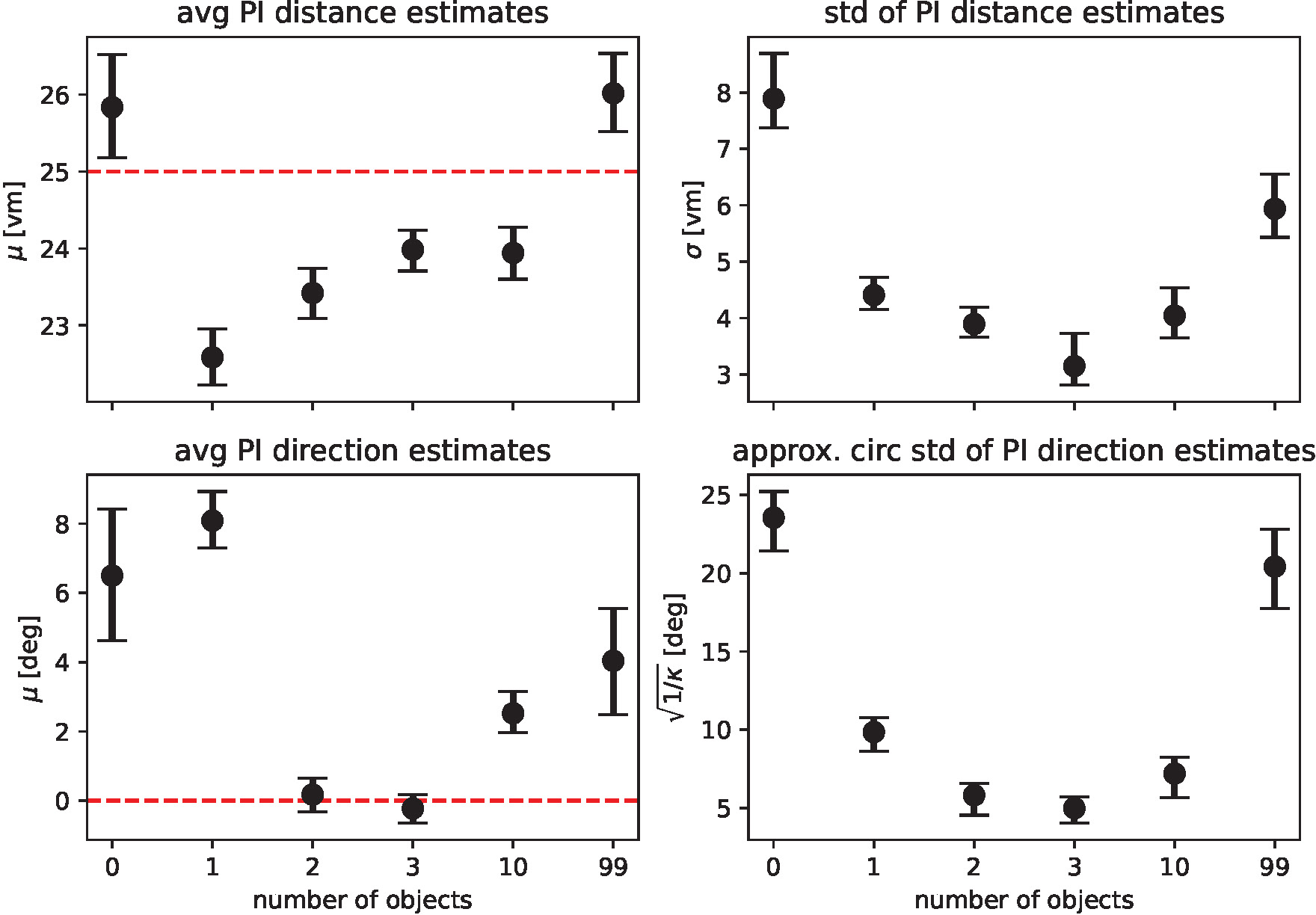
Maximum likelihood parameter estimates for the path integration model. Top row shows maximum likelihood estimates with bootstrapped 95% confidence intervals for the parameters of the distance models fitted on the pooled data of all N=23 participants (*µ* gives the average expected distance, *σ* the standard deviation of expected distances), bottom row shows parameters for the direction models (*µ* gives average expected direction, *κ* is the concentration parameter of the underlying von Mises distribution and has been transformed to circular standard deviation for easier readability). See Eq. 1 and Eq. 2 for a full description of the model functions. Fitted parameters are shown for each experimental condition separately, with overlapping 95% confidence intervals indicating that parameter estimates are similar between conditions. Dashed red lines indicate geometric ground truth where applicable.

## 4 DISCUSSION

In our VR experiment, participants show clear changes in homing performance across environments with varying visual complexity, based on different numbers of objects available. We analysed the average magnitude of homing errors (accuracy), as well as the spread in errors (precision) for each environment. When no objects are present and participants can thus rely on path integration (PI) alone, both accuracy and precision are poor, and individual performance differs considerably. Both measures, accuracy and precision, greatly improve above the level observed for the pure PI condition when one to three objects are added around the goal (Fig. 4).

Going beyond the established configuration of three objects around the goal, this study is the first to our knowledge to also investigate homing in more cluttered environments systematically. For those high clutter environments (namely ten or 99 objects in our experiment), we observe less uniform responses with one share of participants for which both accuracy and precision stay mostly unchanged and another share for which either one or both of these error measures increase considerably, indicating large difficulties to successfully navigate in such cluttered environments (Fig. 4 and Fig. 6). An analysis using linear mixed effects (LME) models, as well as maximum likelihood estimation (MLE) leads us to conclude that homing performance is driven by the number of objects in low clutter environments (zero to three objects), and mainly determined by the individual preferences in cue use of the participants in high clutter environments (three to 99 objects). These individual differences in the influence of environmental complexity on accuracy and precision (Fig. 4 and Fig. 6) are also highlighted by the patterns observed in correlations between accuracy and precision, where higher accuracy is usually associated with higher precision but with variance across participants (Fig. 7).

### 4.1 Comparison to previous studies

Our study was carried out using a desktop-VR setup, so it was of interest to compare the obtained results to those reported previously in similar experiments but with different forms of interaction and presentation (real-world dark-room: Nardini et al. (2008), HMD: Zhao and Warren (2015b); Chen et al. (2017); Sjolund et al. (2018)). In general, we find that variance of position error in the PI condition of our data, when scaling by triangle circumference (*∼*7%) is within the same range as of reported data (Nardini et al. (2008): *∼*10%; Sjolund et al. (2018): *∼*10%; and Chen et al. (2017): *∼*7%). Directional standard deviation in our data is slightly lower but within the same range as in the PI condition and proximal landmark condition as for data reported by Zhao and Warren (2015b) (our data: PI *∼*26*^◦^*, 3 objects *∼*5*^◦^*; Zhao and Warren (2015b): PI *∼*29*^◦^*, combined *∼*10*^◦^*). This indicates that our study broadly captures similar behavioural responses as those reported in previous pertinent studies in the field and that thus our inclusion of high clutter conditions can provide valuable new insights into the individual use of PI and object-based cues. This kind of comparability is especially relevant for studies like those conducted by Kessler et al. (2022, preprint), who aim to develop more comprehensive models of navigation behaviour by using data from different studies.

### 4.2 Low clutter: Path integration and geometric reasoning dominate homing performance

All participants improved in homing performance with up to three objects added around the goal. This can be explained on the basis of object constellation geometry. The first three objects were placed in a triangular arrangement around the goal location. Thus, each additional object further geometrically constrained the region in which the goal could lie. This is because, with only a single object present, the only information available from the object itself is its distance to the goal, which can be memorised by the participant. As predicted by a simple landmark-based homing model by Jetzschke et al. (2017), this leads to a ring-shaped area in which the target could be located based on the object-derived information. We observe goal estimates to be located in this area. As further objects are added, the area becomes more constrained, leading to a fully constrained goal as soon as three objects are present. In other words, to triangulate a specific point on a plane, three distinct references are necessary. Based on this geometric argument, we hypothesised homing performance to increase with each object up to the third one (see “Introduction”). While our findings broadly support this hypothesis, we observe one clear difference to Jetzschke et al. (2017) in the shape of response distributions obtained in our study, which where more spatially constrained. This is unsurprising, since in our version of the task, participants walked around freely in the virtual environment and were not displaced or relocated passively. Therefore, in addition to the object cues, participants could make use of their PI, charged on the outbound path of each trial. Furthermore, the walked triangle itself provides a geometric constraint by directionally limiting the target area based on the approach direction. Thus, the additional information gained via the PI system and the geometry if the outbound path enabled our participants to home more precisely and accurately than if they had access to just the object-derived information in isolation. Overall, for all the low clutter conditions (zero to three objects), the fact that responses were limited to a smaller area than the one observed for the zero object condition indicates that participants combined the different sources of information and that this combination matches the prediction of optimal cue integration in the Bayesian sense, rather than an alternation approach, which would have resulted in a larger spread of responses, rather than a reduced one (Ernst and Banks, 2002; Rohde et al., 2016) (see Fig. 8C-F).

To quantitatively express this combination of spatial cues we combined a model meant to predict distance estimates based on landmarks (Jetzschke et al., 2017) with a model meant to predict directional estimates (Murray and Morgenstern, 2010; Zhao and Warren, 2015b). Together, these two models allowed us to create predictions from both the PI system, as well as any landmark present. We could show that these predictions match the observed spatial pattern of responses, and thus expand on the findings by Jetzschke et al. (2017) for one, two and three landmark objects (see also Fig. 8 middle row). In the condition with one object, this is illustrated by goal estimates being located in only a part of the predicted ring-shaped distribution. In the two object condition, we see a more uniform spread between the predicted two locations, rather than a bimodal distribution of responses, with a clear bias towards the “true” goal location (see Fig. 3 and Fig. 8 middle row). Our MLE models indicate, that this improvement over the “pure” landmark situation described by Jetzschke et al. (2017) can be attributed to the presence of distance information from PI. Finally, with three objects, the goal location is fully spatially constrained and within a Bayesian framework of optimal cue integration, we expect any remaining errors to be attributed to the limitations in cue quality and imprecisions in the encoding of spatial information or execution of movements. Accordingly, we expect a tight distribution of goal estimates around the true goal. For this condition, we actually observe that some of the provided goal estimates are slightly worse than expected for a fully triangulated goal (see Fig. 3 and supplement “S1 Full trajectory figures”). Here again, our MLE models provide additional insight, as we could show that these goal estimates lie within the prediction area of the directional component of our PI model (see Fig. 8). The observed patterns thus indicate that participants integrated object and PI information even when the PI estimate was biased towards underestimating the distance to be travelled.

### 4.3 High clutter: individual preferences in cue use dominate homing performance

For our high clutter conditions (ten and 99 objects) it is hard to disentangle the use of PI and object-based guidance. We cannot simply use the geometric considerations described above, as large inter-individual differences indicate that it is not clear how individuals use the available objects for navigation (Fig. 6).

As outlined by Wolbers and Hegarty (2010) individual navigation abilities might be determined by a variety of factors ranging from how environmental and self-motion cues are perceived and combined, to what cue qualities are personally favoured. Potentially relevant factors may include sex, age, and differences in motion, hippocampal processing, and executive functions (Wolbers and Hegarty, 2010). (Weisberg and Newcombe, 2018) also found evidence that individuals store environmental information in spatial tasks with different degrees of detail and argue for individually different complexities in spatial memory and navigation ability. These individual navigation preferences are supported by the findings of Zanchi et al. (2022) who distinguish two groups of navigators, one group that weights available cues of visual and auditory modality similarly (and combines them optimally in the Bayesian sense), and another group that weights visual cues more strongly. They consider their observation of a preference for visual landmarks in one group to indicate a reset of the directional component of the PI, if visual landmarks are judged to be reliable by the navigator. On a neural level, this reset of PI is facilitated by head direction cells that have been shown to be able to resolve conflicts between different sensory inputs (Zugaro et al., 2003; Valerio and Taube, 2012). Depending on which objects in a condition are considered as reliable cues, different objects might dominate the visual perception of the surroundings for different participants, and if accompanied by high cue weighting, reset PI. Considering that certain trees in our VR are close to the outbound and homing path, possibly a tree’s distance to the first-person avatar and the magnitude of optic flow generated by it, might decide about its dominance in cue integration.

From their experimental results in a cue conflict task between PI and unique landmarks, Zhao and Warren (2015b) suggest that participants use a hybrid model of cue integration. This assessment was later supported by Harootonian et al. (2022) in a cue conflict task with proprioceptive and visual sensory inputs and by Sjolund et al. (2018) who put PI and environmental cues into conflict. In such a hybrid model, body-based and visual cues were combined only when the estimates of PI and landmark guidance were similar. If cues are dissimilar, they will compete for dominance. The switch between combination or competition happens according to the subjectively perceived probability that the respective sensory inputs are correct and refer to the same environmental feature. In our experiment, this is related to the position of goal estimates derived from different cues and the magnitude of difference between them. Interestingly, when PI and landmarks were put into conflicts of intermediate magnitude, (Zhao and Warren, 2015b) and Sjolund et al. (2018) found that homing is dictated by landmark guidance, but with large conflicts people ignore landmarks and use only PI. In the past it has been suggested, that PI might function as a backup system, which is used exclusively when landmark cues are unavailable, unreliable, or in conflict with PI (Cheng et al., 2007). However, in contrast to this idea, Zhao and Warren (2015b) found that participants failed to use PI in trials where landmark cues were unexpectedly removed. When cue competition dominates a in a navigation task, Zhao and Warren (2015b) found indications for cue integration in response variability but at the same time for cue competition in determining response direction, based on the observed variance of responses. They thus concluded that landmarks can reset the orientation of PI without affecting response variance. Using an agent-based dynamic Bayesian actor model, Kessler et al. (2022, preprint) show that an equivalent behaviour can be attributed to multiple noise sources on the levels of perception, motor control, and internal representation, when landmarks reset the internal direction estimate. Also, unlike Zhao and Warren (2015b) suspected, uncertainty in the internal representation is reduced, by preventing it from accumulating additional positional or heading uncertainty in the homing phase.

Our own modelling results, obtained by comparing different variants of an MLE model combining distance and direction information from landmarks and PI, match the conclusions of Kessler et al. (2022, preprint). We find that the standard deviation of both distance and direction estimates is reduced when objects are present (Fig. 9), thus increasing homing precision in multi-cue conditions. Solely for the 99 object condition we find no evidence of increased precision due to the presence of objects. If the cues (PI and objects) were combined optimally in a Bayesian sense, we would not expect the variance to increase in high clutter because more available cues should sharpen the distribution of goal estimates. One possibility to explain this deviation from the initial expectation is a reduction in cue reliability. If landmark or PI-based cues were perceived as less reliable in the high clutter conditions, this would explain the observed absence of a benefit of cue combination. We cannot ultimately determine if the quality of PI or landmark guidance is reduced in clutter, but we argue that with more objects in the scene, the optic flow and therefore the information for PI increases with no identifiable downside. This assumption is further justified by the observation that three participants actually performed most accurately and most precisely in the 99 object condition. For landmarks to aid in reducing uncertainty in the PI system, Kessler et al. (2022, preprint) assume in their model that correspondence between real and perceived landmarks is unambiguously known. Hence, variance reduction requires participants in our experiment to correctly identify objects, a requirement that is not met in the 99 object condition on population level (Fig. 9). Thus, we conclude that when PI and object cues are perceived as conflicting, this does not seem to cause a complete reset of either cue, as we see at least some improvement of precision to the location estimates made by participants. This suggests that, at least for some participants, the reliability of objects in high clutter is reduced.

In addition, it is noteworthy that Kessler et al. (2022, preprint) tested endpoint distributions in different conditions of other studies (Nardini et al., 2008; Zhao and Warren, 2015b; Chen et al., 2017) and found multivariate response normality to be violated, meaning that endpoint distributions deviated from 2D normal distributions, which would be expected if estimates were derived from cues in a Cartesian coordinate system. Our model instead is based on egocentric distance and direction estimates and clearly captures observed curvatures in response clusters (Fig. 8). Thus, we consider an egocentric approach as more reasonable and encourage an explicit investigation of egocentric vs. allocentric encoding of spatial estimates for future studies in the field.

### 4.4 The forest or the trees?

In a visually complex, cluttered environment, people might memorise single objects, constellations of different objects, clearings, or even the edge of the clutter. Therefore, we consider the trees in our experimental design as only “objects” and not “landmarks”, taking into account Montello (2017) who highlights that the term “landmark” is often used in an ambiguous manner.

The quality and reliability of a cue influence its importance and weighting in the process of cue integration. Hence, if objects are considered to be more useful for navigators as a source of spatial information, they should places greater trust in them. If they are correct in their assessment of the usefulness of the object-based cues, this will improve homing performance. We can therefore use the obtained results in our study to draw some inference about the usefulness of the object cues in the different presented environments. Considering the changes in performance across conditions in low and high clutter (see also. 6) in our experiment, we can infer that object guidance is less useful, when no or too many ambiguous objects are available, with one, two, and ten object conditions representing in-between cases. This change in usefulness of the objects is likely caused by processing effort to disambiguate single trees, calculate geometric relations, and encode the goal location from it. Here, the 99 object condition is especially interesting, as participants separate into those who keep good performance and those who get worse, likely based on whether participants are able to disambiguate object cues. We therefore suggest that the 99 object - or high clutter - condition is especially useful for highlighting the differences in the usage of object guidance between individuals. This is supported quantitatively by the fact that comparing the median position error in the three object condition and the 99 object condition on the individual level reveals considerable skew due to a considerable amount of participants in our sample that show distinctly higher median position errors than others (Fig. 6). We thus find a clear group of participants who get lost in the forest, while others seem to manage to stick to their trees.

To further investigate how object cues might have been used, we have employed a series of MLE models of cue combination. While a model variant integrating all available objects is well fitting our empirical data from zero to three objects, we find cue alternation to represent the ten and 99 object conditions better. Note that in this case, the alternation relates to selecting different objects, not to an alternation between PI and object guidance. Nardini et al. (2008) found that adults in their short-range navigation study integrated landmarks and PI in a Bayesian optimal fashion, while children’s behaviour followed a cue alternation model. They conclude that children fail to integrate the two cues. Following the same argumentation, our data suggests that at least some participants in our experiment fail to integrate different objects into one consistent world representation.

We have been able to partially capture the effects of differing abilities of participants to employ objects for guidance by using a cue alternation instead of an integration approach. A model treating every object like one of the three objects immediately surrounding the goal and randomly picking one of them every trial yields at least a rough estimate of population behaviour. While such a model cannot capture when and how individual objects are used as cues, it can be a good first approximation, which further underlines that different participants perceive the objects differently, possibly as individual trees, or as the entirety of a forest.

### 4.5 Path integration and object guidance interplay

Zhao and Warren (2015b), Harootonian et al. (2022), and Kessler et al. (2022, preprint) conclude, that highly reliable cues can reset less reliable ones. For the interpretation our experiment, this might suggest that when the object cues are judged to be highly reliable, they can reset the orientation of PI, while they cannot do so when they are judged to be unreliable. Thus, continued high performance in highly cluttered environments could be the result of successful attempts by the participants to identify specific constellations of objects to arrive at the correct forest clearing. When specific constellations of trees are identified, they result in a reliable object cue that can reset inaccurate PI. Good PI, however, helps to find the correct direction and distance and makes it easier to look for the right object constellations. In contrast, participants who cannot disambiguate between different object constellations might steer towards another, incorrect clearing and suspected goal location, or even disregard all objects and fall back on PI alone. This last strategy should result in a large spread in goal estimates, as can be seen from the zero object condition (Fig. 2 and Fig. 1). Interestingly, from the perspective of cue integration, the interplay of guidance mechanisms (landmarks and PI) might actually lead to worse performance for some participants. If they walk in a wrong direction due to imprecise PI, they might mistake incorrect, but spatially similar, object constellations for their goal location. Such a scenario of mis-identification is inherent to complex cluttered environments and needs to be considered whenever landmarks are not unambiguously identifiable. Studies focusing on the influence of landmark ambiguity in route learning found performance impairment that is correlated with increased activity in the right middle frontal gyrus, indicating additional cognitive demands with each added ambiguity (Janzen and Weststeijn, 2007; Janzen and Jansen, 2010; Strickrodt et al., 2015). This matches the patterns observed in our study and calls into question the assumption made by Kessler et al. (2022, preprint) in their model, where landmarks are always correctly identified. Therefore, application of their generally very attractive model might be limited to use cases with unique landmarks, with low visual complexity, like the analysed datasets from Nardini et al. (2008); Zhao and Warren (2015b); Chen et al. (2017).

As apparent from the comparison between performance accuracy and precision between the zero and 99 object condition for single participants (different coloured dots in Fig. 4), we cannot disentangle if it is PI or landmark guidance that makes participants perform well or badly in high clutter. In other words, people who show good PI in the zero object condition are not necessarily performing well in the 99 object condition, or vice versa. We argue that PI performance in the zero object condition as it stands is not a good predictor for performance in conditions with objects. The MLE models show that PI estimates are updated as objects are added to the environment, as indicated by the change from overshoot to undershoot in the population distance estimates. This kind of qualitative change in the output of the PI system might indicate that the additional spatial information provided by the objects is not only used to gradually improve the estimate provided by the PI system but to actively reset it, as proposed by Zhao and Warren (2015b).

We can also draw conclusions about PI and object guidance and how they might influence each other from the comparison of accuracy and precision in the different conditions. We observe both participants who combine high accuracy in homing with low precision (bottom right quadrants and subfigures a and e in Fig. 7) and end up roughly in the correct location, as well as participants who show low accuracy with high precision (upper left quadrants and subfigures b and c in Fig. 7), who target an incorrect location consistently. Interestingly, low accuracy combined with high precision occurs mainly in conditions with zero, one, and two objects. In the zero object condition, this would imply that the participant’s PI consistently led them to a wrong place. These differences in performance might be related to differential ability to track rotational self-motion, which was demonstrated and linked to differences in cerebellar structure by Chrastil et al. (2017). It also matches observations by Sherrill et al. (2018), who could show individual differences in first-person navigational accuracy, which corresponds to differences in the hippocampus, entorhinal cortex, and thalamus.

When a participant’s PI is repeatedly guiding them into a direction that is not the goal direction, we consider their PI biased (see also population trends in MLE models, Fig. 9). Such a biased PI can also be seen to influence a participant’s goal estimation in other conditions in which objects are present (see Fig. 2a and supplement “S5 PI bias influence on other conditions”). In the one object condition, a biased PI can guide participants to the wrong part of the ring-shaped target area around the only present object (subfigure b in Fig. 7). Thus, combining PI information with object-based cues can cause larger position errors than PI alone, depending on the directional bias of the PI. In the two object condition, the intersection of “rings” around the two objects indicating the correct goal distance leads to two possible goal locations. Here, cue integration with a sufficiently biased PI can lead participants to select the incorrect one repeatedly (subfigure c in Fig. 7). One might also expect inaccurate, but precise participants in the high clutter conditions, as we have just described for the other conditions. However, we do not observe such a pattern for those conditions.

Since one consequence of cue integration is reduced response variance, an alternation from trial to trial between taking different trees into account seems more appropriate in describing behaviour in the high clutter conditions. This implies that participants who struggle in the high clutter conditions, do not confuse single objects, object constellations, or clearings, but rather cannot successfully integrate the information provided by the objects with their PI system and form a consistent image of the forest. Eventually, they end up in a number of wrong places. High accuracy along with low precision can mostly be found in the 99 object condition (bottom right quadrant in Fig. 7). One might expect that participants who show this pattern of performance have problems using the information provided by the objects and fall back on their PI system to solve the task, resulting in error patterns similar to those observed for the zero object condition. However, on closer inspection the participants in question seem to be able to home towards the correct goal with high precision in most trials but target incorrect locations in individual trials, leading to the overall pattern of low precision when analysing the totality of all trials. We see that PI, which is the only cue present in the zero object condition, can also influence target estimation in the other conditions, both positively and negatively. On the one hand, it can help participants select the correct of multiple possible goal locations indicated by objects. On the other hand, a biased PI can also have an opposing effect. Overall, we find that while the PI-driven performance in the zero object condition cannot be used to directly predict performance of participants who get lost in the forest, the spatial patterns of errors we observe and the results of our MLE modelling do indicate a clear interplay between path integration and objects-based cues across all tested conditions.

### 4.6 Conclusion and outlook

In this study, we demonstrate that navigation performance varies in low and high clutter: In low clutter environments, the number of objects and geometric reasoning drive performance, while in high clutter environments, individual differences in cue use are decisive. An MLE model incorporating spatial information from PI and objects reveals that behaviour in low clutter is well described by cue integration. In high clutter, our data reveals that individual participants find different solutions to the task. Some are still able to home accurately and precisely in highly visually complex environments, while others this seems to be impaired in their use of object-based spatial cues. While we are able to outline a variety of reciprocal influences between spatial information from PI and objects, we cannot ultimately elucidate what exact features of objects in clutter of different degrees are used by individual participants, or what exactly determines cue weighting when information is perceived as conflicting.

To further investigate cue weighting of PI guidance and object guidance, our future study designs will incorporate targeted cue conflict trials. Furthermore, to better understand visual processing and the use of object features, we will analyse individual-level eye tracking data. These experimental approaches may be well-suited to reveal why some participants do not see the forest for the trees.

## CONFLICT OF INTEREST STATEMENT

The authors declare that the research was conducted in the absence of any commercial or financial relationships that could be construed as a potential conflict of interest.

## AUTHOR CONTRIBUTIONS

JS: conceptualisation, data curation, formal analysis, methodology, visualisation, writing - original draft, review, and editing. MMM: conceptualisation, data curation, formal analysis, methodology, software, visualisation, supervision, project administration, writing - original draft, review, and editing. PU: investigation, validation, data curation, writing - review, and editing. SM: validation, software, writing - review, and editing. ME, OJNB, NB: funding acquisition, project administration, resources, supervision, writing - review, and editing.

## FUNDING

This work was funded by the Deutsche Forschungsgemeinschaft (DFG) grant 460373158 (https://gepris.dfg.de/gepris/projekt/460373158). OJNB, NB and ME received the funding, MMM, JS and PU were funded as part of the project. We acknowledge the financial support of the Deutsche Forschungsgemeinschaft (DFG) and the Open Access Publication Fund of Bielefeld University for the article processing charge. The funders had no role in study design, data collection and analysis, decision to publish, or preparation of the manuscript.

## ACKNOWLEDGMENTS

We thank Tim Schmoll and the Stats Club at Bielefeld University for valuable input for the statistical analysis of this study. Additionally, we acknowledge that an earlier version of this manuscript was previously shared as a preprint on bioRxiv (Scherer et al., 2023) and we appreciate the valuable feedback received during the preprint stage. Moreover, we recognize the prior publication of this work as a PhD thesis titled “Finding Back Home: From Vector Models to Virtual Forests” by Martin M. Müller (Müller, 2023).

## SUPPLEMENTAL DATA

see attached

## DATA AVAILABILITY STATEMENT

The study dataset that supports the findings of this study along with Jupyter Notebooks containing the full analysis pipeline are openly available. Data is available at https://pub.uni-bielefeld.de/record/2982046. Analysis code with accompanying instructions for execution is available on GitLab at https://gitlab.ub.uni-bielefeld.de/jonas.scherer/not_seeing_the_forest_for_the_trees_combination_of_path_integration_and_landmark_cues_in_human_virtual_navigation. Implementation details of the Unity3D project and an executable Unity project of this study’s task are available on GitLab at https://gitlab.ub.uni-bielefeld.de/virtual_navigation_tools/unity_vnt_showcase_triangle_completion.

## Footnotes

1 https://docs.python.org/3.9/reference/index.html

2 https://stats.oarc.ucla.edu/r/library/r-library-contrast-coding-systems-for-categorical-variables/

## Notes

### Competing Interest Statement

The authors have declared no competing interest.

### Summary of Updates

This updated version of the manuscript clarifies thoughts in the introduction and the discussion. More published studies are taken into consideration. We added a subfigure to figure 2, and two figures to the supplementary material. The Methods section was moved up after the introduction.

## REFERENCES

Alais, D. and Burr, D. (2019). Cue combination within a bayesian framework. Multisensory processes: The auditory perspective, 9–31

Chen, X., McNamara, T. P., Kelly, J. W., and Wolbers, T. (2017). Cue combination in human spatial navigation. Cognitive Psychology 95, 105–144

Cheng, K., Shettleworth, S. J., Huttenlocher, J., and Rieser, J. J. (2007). Bayesian integration of spatial information. Psychological bulletin 133, 625

Chrastil, E. R., Nicora, G. L., and Huang, A. (2019). Vision and proprioception make equal contributions to path integration in a novel homing task. Cognition 192, 103998

Chrastil, E. R., Sherrill, K. R., Aselcioglu, I., Hasselmo, M. E., and Stern, C. E. (2017). Individual differences in human path integration abilities correlate with gray matter volume in retrosplenial cortex, hippocampus, and medial prefrontal cortex. Eneuro 4

Chrastil, E. R. and Warren, W. H. (2017). Rotational error in path integration: encoding and execution errors in angle reproduction. Experimental brain research 235, 1885–1897

Chrastil, E. R. and Warren, W. H. (2021). Executing the homebound path is a major source of error in homing by path integration. Journal of Experimental Psychology: Human Perception and Performance 47, 13

Ernst, M. O. and Banks, M. S. (2002). Humans integrate visual and haptic information in a statistically optimal fashion. Nature 415, 429–433

Etienne, A. S. and Jeffery, K. J. (2004). Path integration in mammals. Hippocampus 14, 180–192

Faul, F., Erdfelder, E., Lang, A.-G., and Buchner, A. (2007). G* power 3: A flexible statistical power analysis program for the social, behavioral, and biomedical sciences. Behavior research methods 39, 175–191

Glasauer, S., Amorim, M. A., Viaud-Delmon, I., and Berthoz, A. (2002). Differential effects of labyrinthine dysfunction on distance and direction during blindfolded walking of a triangular path. Experimental Brain Research 145, 489–497

Glasauer, S. and Shi, Z. (2022). Individual beliefs about temporal continuity explain variation of perceptual biases. Scientific Reports 12, 10746

Goeke, C. M., Planera, S., Finger, H., and Kö nig, P. (2016). Bayesian alternation during tactile augmentation. Frontiers in Behavioral Neuroscience 10. doi:10.3389/fnbeh.2016.00187

Harootonian, S. K., Ekstrom, A. D., and Wilson, R. C. (2022). Combination and competition between path integration and landmark navigation in the estimation of heading direction. PLoS computational biology 18, e1009222

Harootonian, S. K., Wilson, R. C., Hejtmánek, L., Ziskin, E. M., and Ekstrom, A. D. (2020). Path integration in large-scale space and with novel geometries: Comparing vector addition and encoding-error models. PLoS computational biology 16, e1007489

Hoinville, T. and Wehner, R. (2018). Optimal multiguidance integration in insect navigation. Proceedings of the National Academy of Sciences 115, 2824–2829

Janzen, G. and Jansen, C. (2010). A neural wayfinding mechanism adjusts for ambiguous landmark information. NeuroImage 52, 364–370

Janzen, G. and Weststeijn, C. G. (2007). Neural representation of object location and route direction: an event-related fmri study. Brain research 1165, 116–125

Jetzschke, S., Ernst, M. O., Froehlich, J., and Boeddeker, N. (2017). Finding home: Landmark ambiguity in human navigation. Frontiers in behavioral neuroscience 11, 132

Jetzschke, S., Ernst, M. O., Moscatelli, A., and Boeddeker, N. (2016). Going round the bend: Persistent personal biases in walked angles. Neuroscience letters 617, 72–75

Johnson, P. C. (2014). Extension of nakagawa & schielzeth’s r2glmm to random slopes models. Methods in ecology and evolution 5, 944–946

Kearns, M. J., Warren, W. H., Duchon, A. P., and Tarr, M. J. (2002). Path integration from optic flow and body senses in a homing task. Perception 31, 349–374

Kessler, F., Frankenstein, J., and Rothkopf, C. A. (2022, preprint). A dynamic bayesian actor model explains endpoint variability in homing tasks. bioRxiv, 2022–11

Loomis, J. M., Klatzky, R. L., Golledge, R. G., Cicinelli, J. G., Pellegrino, J. W., and Fry, P. A. (1993). Nonvisual navigation by blind and sighted: assessment of path integration ability. Journal of Experimental Psychology: General 122, 73

Mallot, H. A. and Lancier, S. (2018). Place recognition from distant landmarks: human performance and maximum likelihood model. Biological cybernetics 112, 291–303

McNamara, T. P. and Chen, X. (2022). Bayesian decision theory and navigation. Psychonomic Bulletin & Review, 1–32

Montello, D. R. (2017). Landmarks are exaggerated. KI-Künstliche Intelligenz 31, 193–197

Müller, M. (2023). Finding back home: from vector models to virtual forests. Ph.D. thesis, Bielefeld University

Murray, R. F. and Morgenstern, Y. (2010). Cue combination on the circle and the sphere. Journal of vision 10, 15–15

Müller, M., Scherer, J., Unterbrink, P., Bertrand, O. J. N., Egelhaaf, M., and Boeddeker, N. (2023). The virtual navigation toolbox: Providing tools for virtualnavigation experiments. PLoS One Forthcoming

Nakagawa, S. and Schielzeth, H. (2013). A general and simple method for obtaining r2 from generalized linear mixed-effects models. Methods in ecology and evolution 4, 133–142

Nardini, M., Jones, P., Bedford, R., and Braddick, O. (2008). Development of cue integration in human navigation. Current Biology 18, 689–693. doi:10.1016/j.cub.2008.04.021

R Core Team (2022). R: A Language and Environment for Statistical Computing. R Foundation for Statistical Computing, Vienna, Austria

Rohde, M., van Dam, L. C., and Ernst, M. O. (2016). Statistically optimal multisensory cue integration: A practical tutorial. Multisensory research 29, 279–317

Roy, C., Wiebusch, D., Botsch, M., and Ernst, M. O. (2023). Did it move? humans use spatio-temporal landmark permanency efficiently for navigation. Journal of Experimental Psychology: General 152, 448

Scherer, J., Müller, M. M., Unterbrink, P., Meier, S., Egelhaaf, M., Bertrand, O. J. N., et al. (2023). Not seeing the forest for the trees: Combination of path integration and landmark cues in human virtual navigation. bioRxiv doi:10.1101/2023.10.25.563902

Schielzeth, H., Dingemanse, N. J., Nakagawa, S., Westneat, D. F., Allegue, H., Teplitsky, C., et al. (2020). Robustness of linear mixed-effects models to violations of distributional assumptions. Methods in ecology and evolution 11, 1141–1152

Sherrill, K. R., Chrastil, E. R., Aselcioglu, I., Hasselmo, M. E., and Stern, C. E. (2018). Structural differences in hippocampal and entorhinal gray matter volume support individual differences in first person navigational ability. Neuroscience 380, 123–131

Sjolund, L. A., Kelly, J. W., and McNamara, T. P. (2018). Optimal combination of environmental cues and path integration during navigation. Memory & Cognition 46, 89–99

Strickrodt, M., O’Malley, M., and Wiener, J. M. (2015). This place looks familiar—how navigators distinguish places with ambiguous landmark objects when learning novel routes. Frontiers in Psychology 6, 1936

UCLA Statistical Consulting Group (2021). R library contrast coding systems for categorical variables. https://stats.oarc.ucla.edu/r/library/r-library-contrast-coding-systems-for-categorical-variables/ Accessed: 2024-29-01

Valerio, S. and Taube, J. S. (2012). Path integration: how the head direction signal maintains and corrects spatial orientation. Nature neuroscience 15, 1445–1453

Van Rossum, G. and Drake, F. L. (2009). Python 3 Reference Manual (Scotts Valley, CA: CreateSpace)

Walter, J. L., Essmann, L., König, S. U., and Kö nig, P. (2022). Finding landmarks-an investigation of viewing behavior during spatial navigation in vr using a graph-theoretical analysis approach. PLoS Computational Biology 18, e1009485

Weisberg, S. M. and Newcombe, N. S. (2018). Cognitive maps: Some people make them, some people struggle. Current directions in psychological science 27, 220–226

Widdowson, C. and Wang, R. F. (2022). Human navigation in curved spaces. Cognition 218, 104923

Wiener, J. M. and Mallot, H. A. (2006). Path complexity does not impair visual path integration. Spatial cognition and computation 6, 333–346

Wolbers, T. and Hegarty, M. (2010). What determines our navigational abilities? Trends in cognitive sciences 14, 138–146

Wolbers, T., Wiener, J. M., Mallot, H. A., and Büchel, C. (2007). Differential recruitment of the hippocampus, medial prefrontal cortex, and the human motion complex during path integration in humans. Journal of Neuroscience 27, 9408–9416

Wozny, D. R., Beierholm, U. R., and Shams, L. (2010). Probability matching as a computational strategy used in perception. PLoS computational biology 6, e1000871

Xie, Y., Bigelow, R. T., Frankenthaler, S. F., Studenski, S. A., Moffat, S. D., and Agrawal, Y. (2017). Vestibular loss in older adults is associated with impaired spatial navigation: data from the triangle completion task. Frontiers in neurology 8, 173

Zanchi, S., Cuturi, L. F., Sandini, G., and Gori, M. (2022). Interindividual differences influence multisensory processing during spatial navigation. Journal of Experimental Psychology: Human Perception and Performance 48, 174

Zhao, M. and Warren, W. H. (2015a). Environmental stability modulates the role of path integration in human navigation. Cognition 142, 96–109

Zhao, M. and Warren, W. H. (2015b). How you get there from here: Interaction of visual landmarks and path integration in human navigation. Psychological science 26, 915–924

Zhao, M. and Warren, W. H. (2018). Non-optimal perceptual decision in human navigation. Behavioral and Brain Sciences 41

Zugaro, M. B., Arleo, A., Berthoz, A., and Wiener, S. I. (2003). Rapid spatial reorientation and head direction cells. Journal of Neuroscience 23, 3478–3482

