## Supplementary Material for "Not seeing the forest for the trees: Combination of path integration and landmark cues in human virtual navigation"

### 1 SUPPLEMENTARY MATERIAL

#### *S1 Full trajectory figures.*

Here we provide figures showing full trajectory data that depict all trials from Figure 3. Most trajectories are straight when no or only few objects are present but for some participants they become more complex and curved with ten and 99 objects.

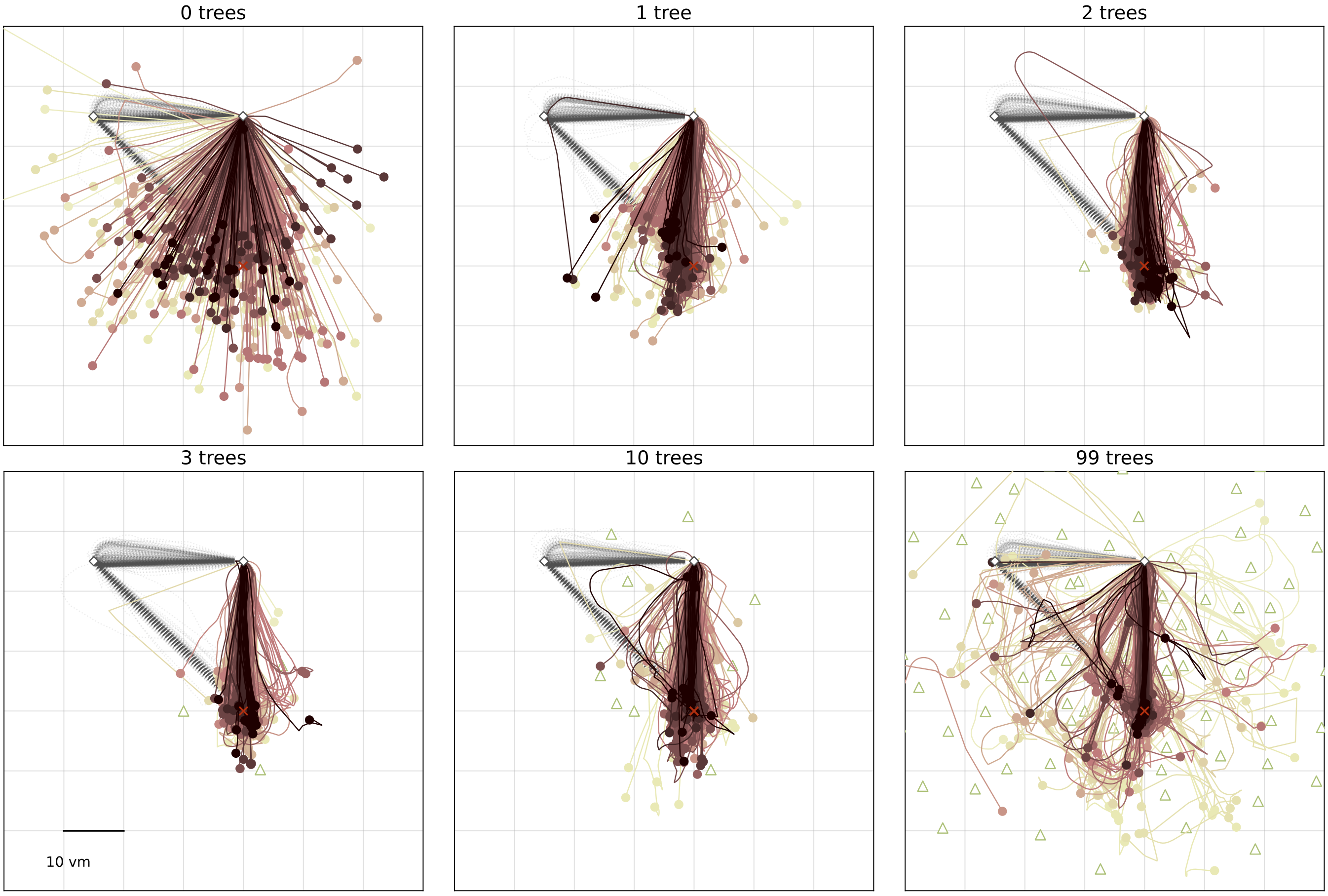


**Figure S1.** Trajectory endpoints of single trials are depicted as dots. Colours marking individual Participants are matched across all figures, based on a given participant's median position error in the 99-object condition (refer to Fig. 4 top right). Marker size does not match actual tree dimensions. Cutout does not show all objects, please see Fig. 1 for a more zoomed-out view of the environment.

#### *S2 Session order effect.*

In our analysis, we observe a trend of reduced position error over sessions (Fig. S2). This is unsurprising, as placement of objects and shape of the outbound path do not vary across trials and conditions. Solely the number of objects from a given set of objects with fixed positions differs between conditions. However, including session number as an additional fixed effect in a regression model results in only slight improvement of fit for accuracy (without session: AIC≈1163.4, BIC≈1193.6; with session: AIC≈1150.9, BIC≈1194; likelihood ratio test: $\chi^2$ ≈18.58, logLik1=-574.7, logLik2=565.4, p≈0.00033) and for precision (without session: AIC≈1385.8, BIC≈1416; with session: AIC≈1346.6, BIC≈1389.7; likelihood ratio test: $\chi^2$ ≈45.18, logLik1=-685.9, logLik2=663.3, p<0.0001). As the order effect is small and not the primary focus of this study we do not consider it in further analyses. In our linear regression analysis, we average across trials and sessions using median position errors and position error standard deviation (scatter plot in Fig. 4). We also provide a linear mixed effects model on single-trial data (supplement Tab. S1). When comparing the differences between the zero and three object condition and between the three and 99 object condition for session 1 and session 4, we do not find a significant change between the sessions (Fig. S3) in accuracy (Wilcoxon signed-rank test: 0 vs 3 objects: Z=107, p≈0.36, d≈0.26; 3 vs 99 objects: Z=95, p≈0.2, d≈0.2)). Only in precision session 4 shows significantly lower values than session 1 (Wilcoxon signed-rank test: 0 vs 3 objects: Z=129, p≈0.8, d≈0.05; 3 vs 99 objects: Z=60, p≈0.016, d≈0.76)).

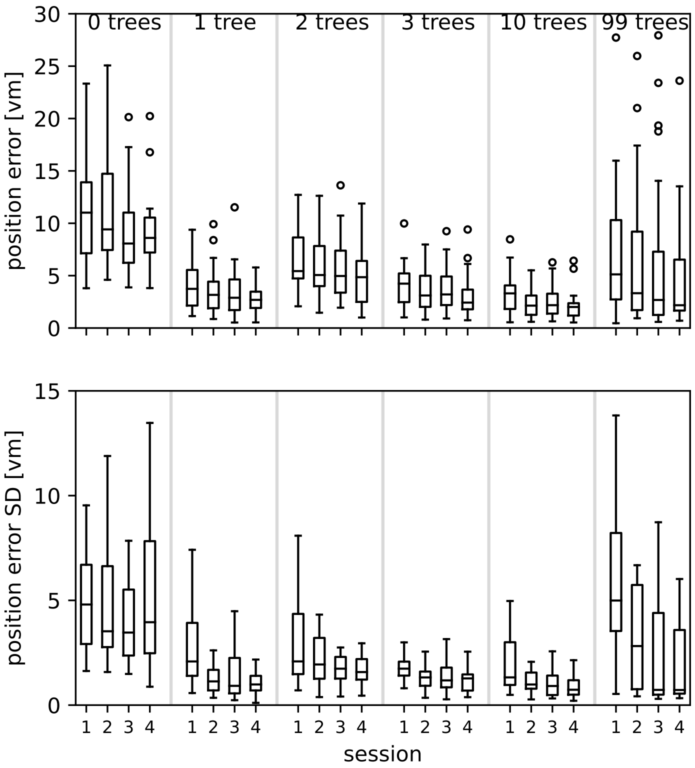


**Figure S2.** Order effect within the different conditions on position error accuracy (top) and precision (bottom). N=23 participants completed four sessions with six repetitions (n=24) in each condition.

**Table S1. Performance of linear regression models on single trial basis**. The model predicts the position error from the condition (number of objects). In all models condition is ordinal and polynomials from 1st to 5th degree are fitted. est.: estimate; s.e.: standard error; t: t value; d.f.:degrees of freedom; p: p-value

$model : \ln(position\ error) \sim condition\ ordinal$
$AIC = 7893.5, BIC = 7942.2$

| | est. | s.e. | t | d.f. | p |
|---|---|---|---|---|---|
| intercept | 1.3 | 0.08 | 15.61 | 22 | <0.001 |
| condition $^1$ | -0.77 | 0.03 | -22.69 | 3243.05 | <0.001 |
| condition $^2$ | 0.85 | 0.03 | 25.06 | 3243.05 | <0.001 |
| condition $^3$ | 0.04 | 0.03 | 1.1 | 3243.05 | 0.27 |
| condition $^4$ | -0.14 | 0.03 | -4.03 | 3243.05 | <0.001 |
| condition $^5$ | -0.10 | 0.03 | -2.94 | 3243.05 | <0.001 |

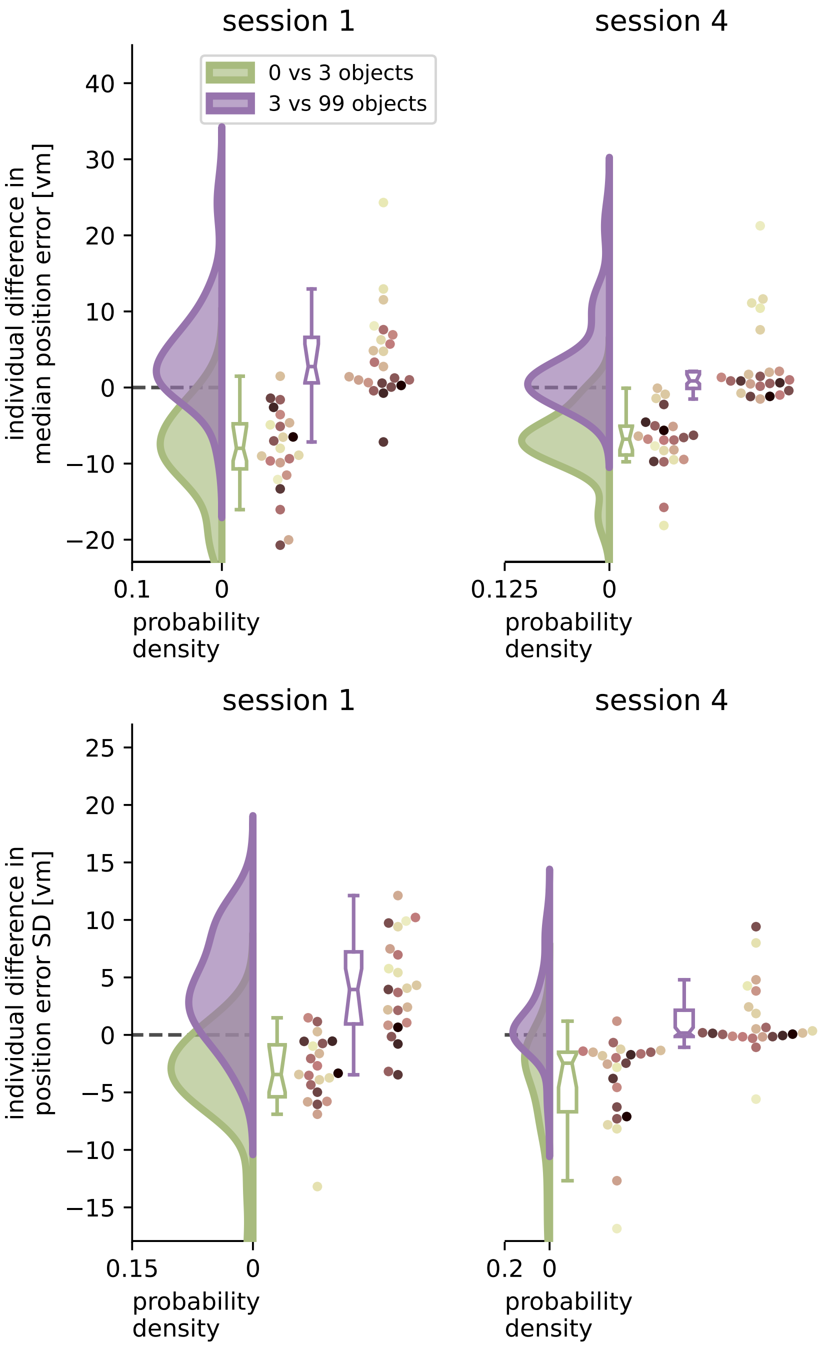


**Figure S3.** Performance improvement in accuracy and precision of position error from zero to three objects and performance impairment from three to 99 objects separated for session 1 and session 4. Green and purple curves indicate kernel-density estimations fitted to the difference in median position error (left) or position error standard deviation (SD) (right) between zero and three objects (green), or three and 99 objects (purple), respectively, of N=23 single participants with n=24 repetitions within each condition (see dots). Boxplot notches indicate 95% confidence intervals around the median. Colours marking individual Participants are matched across all figures, based on a given participant's median position error in the 99-object condition (refer to Fig. 4 top right).

##### *S3 Separating effects of low and high clutter.*

We present two approaches, one based on a quadratic regression and one based on forward difference contrast coding, to justify splitting the dataset into two separate effects, an effect of low clutter and an effect of high clutter.

###### *1) Linear vs quadratic fit*

We observe a significant linear and a significant quadratic component in our linear mixed effects regression models for both accuracy and precision (see Results section ”Disentangling the effect of different degrees of clutter”). A simple linear fit does not well reflect the visually observed pattern of position error decrease from zero to three objects and increase from three to 99 objects (see Fig. 4 and 5). The significant quadratic component matches the observation better, which is confirmed by a comparison of the linear and the quadratic component of our linear mixed models above. The comparison shows the quadratic function yielding a better fit to our data (for accuracy: linear: AIC≈293, BIC≈301.8; quadratic: AIC≈255.4, BIC≈267.1; likelihood ratio test: $\chi^2$ ≈39.63, logLik1=-143.5, logLik2=123.7, p<0.0001; and for precision AIC≈297.7, BIC≈306.5; quadratic: AIC≈241.5, BIC≈253.2; likelihood ratio test: $\chi^2$ ≈58.23, logLik1=-145.8, logLik2=-116.7, p<0.0001) (see Tab. 1). An evaluation of the quadratic fits reveals theoretical minima for position error (3.19) and its standard deviation (2.66) close to the three object condition, identifying this condition as the one with the smallest homing errors (see Fig. 5). This speaks in favour of two separate effects with an error decrease for low clutter conditions (zero to three objects), and an error increase for high clutter conditions (three to 99 objects).

###### *2) Forward difference contrast coding*

Alternatively to analysing the linear and quadratic components of our linear mixed model, we can re-state the model using forward difference contrast coding. In forward difference contrast coding, “the mean of the dependent variable for one level of the categorical variable is compared to the mean of the dependent variable for the next level” (UCLA Statistical Consulting Group, 2021). The underlying coding matrix allows for comparing the central tendency of the dependent variable of each class with the subsequent class on the axis of the independent variable. In our case, this means that the zero object condition is compared to the one object condition, which in turn is compared to the two object condition, and so on. For our data, this approach results in significant negative slopes for the conditions with one, two and three objects, and in significant positive slopes for ten and 99 objects, for both accuracy (Tab. 1 model 3) and precision (Tab. 1 model 6). Note that for precision the estimate from the two to three object condition shows only a trend, indicating no significant improvement in precision. Again, this analysis speaks in favour of two separate effects for low clutter and high clutter conditions.

##### *S4 MLE model summaries.*

Here we provide detailed comparisons of the fitted MLE models separately for each condition (Tab. S2-S7)

**Table S2.** Summary table of MLE model fits for the zero object condition. For an overview of model components see Table 2. Model fits within each subdivision can be compared directly, since they belong the the same model family.

| model ID | log Likelihood | BIC | AIC |
|---|---|---|---|
| $g^*_{PI}$ | -1902.68 | 3817.96 | 3809.35 |
| $f^{flat}_{PI}$ | -1003.48 | 2019.57 | 2010.96 |
| $f^*_{PI}$ | -304.07 | 620.75 | 612.15 |
| $g^*_{PI} \times f^*_{PI}$ | -2206.75 | 4438.71 | 4421.50 |
| $g^*_{PI} \times f^{flat}_{PI}$ | -2906.16 | 5837.52 | 5820.31 |

**Table S3.** Summary table of MLE model fits for the one object condition. For an overview of model components see Table 2. Model fits within each subdivision can be compared directly, since they belong the the same model family.

| model ID | log Likelihood | BIC | AIC |
|---|---|---|---|
| $g^{*\,0objs}_{PI}$ | -1767.60 | 3547.81 | 3539.20 |
| $g^*_{PI}$ | -1590.35 | 3193.31 | 3184.69 |
| $f^{flat}_{PI}$ | -1007.16 | 2026.93 | 2018.31 |
| $f^{*\,0objs}_{PI}$ | -78.40 | 169.41 | 160.80 |
| $f^*_{PI}$ | 185.07 | -357.53 | -366.14 |
| $\prod g^{true-dist}_{LM}$ | -1344.11 | 2700.82 | 2692.21 |
| $g^*_{PI} \times f^*_{PI} \times \prod g^{true-dist}_{LM}$ | -2749.38 | 5536.60 | 5510.92 |
| $g^*_{PI} \times f^{*\,0objs}_{PI} \times \prod g^{true-dist}_{LM}$ | -3012.85 | 6063.55 | 6037.86 |
| $g^*_{PI} \times f^{flat}_{PI} \times \prod g^{true-dist}_{LM}$ | -3941.61 | 7921.06 | 7895.37 |
| $g^{*\,0objs}_{PI} \times f^*_{PI} \times \prod g^{true-dist}_{LM}$ | -2926.63 | 5891.10 | 5865.42 |
| $g^{*\,0objs}_{PI} \times f^{*\,0objs}_{PI} \times \prod g^{true-dist}_{LM}$ | -3190.11 | 6418.05 | 6392.37 |
| $g^{*\,0objs}_{PI} \times f^{flat}_{PI} \times \prod g^{true-dist}_{LM}$ | -4118.86 | 8275.56 | 8249.88 |

**Table S4.** Summary table of MLE model fits for the two object condition. For an overview of model components see Table 2. Model fits within each subdivision can be compared directly, since they belong the the same model family.

| model ID | log Likelihood | BIC | AIC |
|---|---|---|---|
| $g_{PI}^{*\,0objs}$ | -1731.15 | 3474.92 | 3466.31 |
| $g_{PI}^{*}$ | -1525.18 | 3062.99 | 3054.37 |
| $f_{PI}^{flat}$ | -1008.99 | 2030.61 | 2021.99 |
| $f_{PI}^{*\,0objs}$ | -65.62 | 143.85 | 135.23 |
| $f_{PI}^{*}$ | 476.24 | -939.85 | -948.47 |
| $\prod g_{LM}^{true-dist}$ | -2612.62 | 5250.48 | 5233.25 |
| $g_{PI}^{*} \times f_{PI}^{*} \times \prod g_{LM}^{true-dist}$ | -3661.57 | 7373.61 | 7339.41 |
| $g_{PI}^{*} \times f_{PI}^{*\,0objs} \times \prod g_{LM}^{true-dist}$ | -4203.42 | 8457.31 | 8423.11 |
| $g_{PI}^{*} \times f_{PI}^{flat} \times \prod g_{LM}^{true-dist}$ | -5146.80 | 10344.07 | 10309.87 |
| $g_{PI}^{*\,0objs} \times f_{PI}^{*} \times \prod g_{LM}^{true-dist}$ | -3867.54 | 7785.55 | 7751.35 |
| $g_{PI}^{*\,0objs} \times f_{PI}^{*\,0objs} \times \prod g_{LM}^{true-dist}$ | -4409.39 | 8869.25 | 8835.05 |
| $g_{PI}^{*\,0objs} \times f_{PI}^{flat} \times \prod g_{LM}^{true-dist}$ | -5352.77 | 10756.00 | 10721.81 |

**Table S5.** Summary table of MLE model fits for the three object condition. For an overview of model components see Table 2. Model fits within each subdivision can be compared directly, since they belong the the same model family.

| model ID | log Likelihood | BIC | AIC |
|---|---|---|---|
| $g_{PI}^{*\,0objs}$ | -1691.17 | 3394.95 | 3386.35 |
| $g_{PI}^{*}$ | -1403.12 | 2818.84 | 2810.23 |
| $f_{PI}^{flat}$ | -1005.32 | 2023.25 | 2014.64 |
| $f_{PI}^{*\,0objs}$ | -63.55 | 139.70 | 131.09 |
| $f_{PI}^{*}$ | 559.06 | -1105.50 | -1114.11 |
| $\prod g_{LM}^{true-dist}$ | -4033.60 | 8105.04 | 8079.37 |
| $g_{PI}^{*} \times f_{PI}^{*} \times \prod g_{LM}^{true-dist}$ | -4877.66 | 9818.37 | 9775.74 |
| $g_{PI}^{*} \times f_{PI}^{*\,0objs} \times \prod g_{LM}^{true-dist}$ | -5500.27 | 11063.58 | 11020.95 |
| $g_{PI}^{*} \times f_{PI}^{flat} \times \prod g_{LM}^{true-dist}$ | -6442.04 | 12947.12 | 12904.49 |
| $g_{PI}^{*\,0objs} \times f_{PI}^{*} \times \prod g_{LM}^{true-dist}$ | -5165.72 | 10394.49 | 10351.85 |
| $g_{PI}^{*\,0objs} \times f_{PI}^{*\,0objs} \times \prod g_{LM}^{true-dist}$ | -5788.32 | 11639.69 | 11597.06 |
| $g_{PI}^{*\,0objs} \times f_{PI}^{flat} \times \prod g_{LM}^{true-dist}$ | -6730.10 | 13523.24 | 13480.60 |

**Table S6.** Summary table of MLE model fits for the ten object condition. For an overview of model components see Table 2. Model fits within each subdivision can be compared directly, since they belong the the same model family.

| model ID | log Likelihood | BIC | AIC |
|---|---|---|---|
| $g_{PI}^{*0objs}$ | -1732.74 | 3478.10 | 3469.48 |
| $g_{PI}^{*}$ | -1551.43 | 3115.49 | 3106.86 |
| $f_{PI}^{flat}$ | -1012.67 | 2037.96 | 2029.34 |
| $f_{PI}^{*0objs}$ | -62.93 | 138.48 | 129.85 |
| $f_{PI}^{*}$ | 359.59 | -706.56 | -715.18 |
| $\prod g_{LM}^{true-dist}$ | -41593.09 | 83312.42 | 83227.77 |
| $\sum g_{LM}^{true-dist}$ | -423.04 | 972.32 | 887.67 |
| $g_{PI}^{*} \times f_{PI}^{*} \times \sum g_{LM}^{true-dist}$ | -1614.88 | 3381.25 | 3280.05 |
| $g_{PI}^{*} \times f_{PI}^{*0objs} \times \sum g_{LM}^{true-dist}$ | -2037.40 | 4226.28 | 4125.08 |
| $g_{PI}^{*} \times f_{PI}^{flat} \times \sum g_{LM}^{true-dist}$ | -2987.15 | 6125.77 | 6024.57 |
| $g_{PI}^{*0objs} \times f_{PI}^{*} \times \sum g_{LM}^{true-dist}$ | -1796.19 | 3743.86 | 3642.66 |
| $g_{PI}^{*0objs} \times f_{PI}^{*0objs} \times \sum g_{LM}^{true-dist}$ | -2218.71 | 4588.90 | 4487.70 |
| $g_{PI}^{*0objs} \times f_{PI}^{flat} \times \sum g_{LM}^{true-dist}$ | -3168.45 | 6488.38 | 6387.18 |

**Table S7.** Summary table of MLE model fits for the 99 object condition. For an overview of model components see Table 2. Model fits within each subdivision can be compared directly, since they belong the the same model family.

| model ID | log Likelihood | BIC | AIC |
|---|---|---|---|
| $g_{PI}^{*\,0objs}$ | -1732.01 | 3476.57 | 3468.02 |
| $g_{PI}^{*}$ | -1696.02 | 3404.59 | 3396.04 |
| $f_{PI}^{flat}$ | -974.07 | 1960.70 | 1952.15 |
| $f_{PI}^{*\,0objs}$ | -231.06 | 474.66 | 466.12 |
| $f_{PI}^{*}$ | -215.49 | 443.53 | 434.99 |
| $\prod g_{LM}^{true-dist}$ | -inf | inf | inf |
| $\sum g_{LM}^{true-dist}$ | 56.78 | 1128.46 | 520.51 |
| $g_{PI}^{*} \times f_{PI}^{*} \times \sum g_{LM}^{true-dist}$ | -1854.73 | 4976.59 | 4364.27 |
| $g_{PI}^{*} \times f_{PI}^{*\,0objs} \times \sum g_{LM}^{true-dist}$ | -1870.30 | 5007.71 | 4395.39 |
| $g_{PI}^{*} \times f_{PI}^{flat} \times \sum g_{LM}^{true-dist}$ | -2613.31 | 6493.75 | 5881.43 |
| $g_{PI}^{*\,0objs} \times f_{PI}^{*} \times \sum g_{LM}^{true-dist}$ | -1890.72 | 5048.56 | 4436.24 |
| $g_{PI}^{*\,0objs} \times f_{PI}^{*\,0objs} \times \sum g_{LM}^{true-dist}$ | -1906.29 | 5079.69 | 4467.37 |
| $g_{PI}^{*\,0objs} \times f_{PI}^{flat} \times \sum g_{LM}^{true-dist}$ | -2649.30 | 6565.73 | 5953.41 |

#### *S5 PI bias influence on other conditions.*

When a participant's PI is repeatedly guiding them into a direction that is not the goal direction, we consider their PI biased (see also population trends in MLE models, Fig. 9). Such a biased PI can also be seen to influence a participant's goal estimation in other conditions in which objects are present. Below, we show the bootstrapped parameters of our path integration model for one such participant (Fig. S4) as an example. Data of the same participant is also plotted in Fig. 2a. The model parameters show that the biased PI observed in the zero object condition also guides the participant in the same wrong direction in the 99 object condition. The 95% confidence intervals of the direction and the distance estimate of these two conditions overlap, indicating a return of the bias present in the zero object condition.

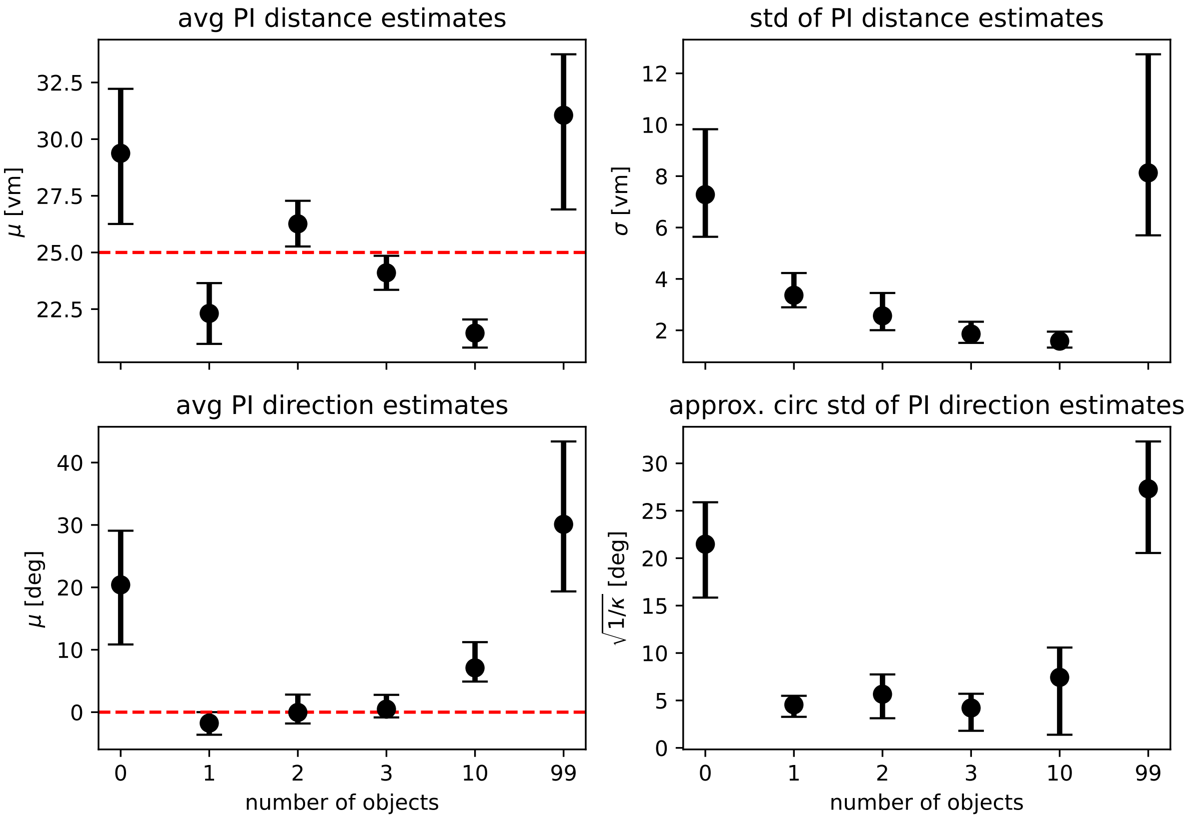


**Figure S4.** Maximum likelihood parameter estimates for the path integration model of one single participant also plotted in Fig. 2a. Top row shows maximum likelihood estimates with bootstrapped 95% confidence intervals for the parameters of the distance models ($\mu$ gives the average expected distance, $\sigma$ the standard deviation of expected distances), bottom row shows parameters for the direction models ($\mu$ gives average expected direction, $\kappa$ is the concentration parameter of the underlying von Mises distribution and has been transformed to circular standard deviation for easier readability). See Eq. 1 and Eq. 2 for a full description of the model functions. Fitted parameters are shown for each experimental condition separately, with overlapping 95% confidence intervals indicating that parameter estimates are similar between conditions. Dashed red lines indicate geometric ground truth where applicable.
